# CRTC-1 balances histone trimethylation and acetylation to promote longevity

**DOI:** 10.1101/2022.08.31.506037

**Authors:** Carlos G. Silva-García, Laura I. Láscarez-Lagunas, Katharina Papsdorf, Caroline Heintz, Aditi Prabhakar, Christopher S. Morrow, Lourdes Pajuelo Torres, Arpit Sharma, Jihe Liu, Monica P. Colaiácovo, Anne Brunet, William B. Mair

## Abstract

Loss of function during ageing is accompanied by transcriptional drift, altering gene expression and contributing to a variety of age-related diseases. CREB-regulated transcriptional coactivators (CRTCs) have emerged as key regulators of gene expression that might be targeted to promote longevity. Here, we define the role of the *Caenorhabditis elegans* CRTC-1 in the epigenetic regulation of longevity. Endogenous CRTC-1 binds chromatin factors, including components of the COMPASS complex, which trimethylates lysine 4 on histone H3 (H3K4me3). CRISPR editing of endogenous CRTC-1 reveals that the CREB-binding domain in neurons is specifically required for H3K4me3-dependent longevity. However, this effect is independent of CREB but instead acts via the transcription factor AP-1. Strikingly, CRTC-1 also mediates global histone acetylation levels, and this acetylation is essential for H3K4me3-dependent longevity. Indeed, overexpression of an acetyltransferase enzyme is sufficient to promote longevity in wild-type worms. CRTCs, therefore, link energetics to longevity by critically fine-tuning histone acetylation and methylation to promote healthy ageing.

## MAIN TEXT

Modulation of transcriptional regulators has emerged as an evolutionarily conserved mechanism to slow ageing and promote longevity^1^. Gene expression can be optimised for health and longevity by manipulating transcription factors and their regulators, making them attractive targets to treat or prevent age-related diseases. Recent studies in the nematode *C. elegans* have highlighted a family of cofactors, cAMP response element-binding protein (CREB)-regulated transcriptional coactivators (CRTCs), as novel modulators of ageing that link energy sensing to transcription^2,3^. Although CRTCs are traditionally described as transcriptional coactivators, they are in fact multifunctional. CRTCs bind additional basic leucine zipper (bZIP) transcription factors than CREB, and beyond transcription, directly regulate other cellular processes including endoplasmic reticulum (ER)- to-Golgi transport via coat protein complex II (COPII)-mediated vesicle trafficking, RNA splicing and chromatin regulation^4–7^. However, despite their importance in multiple age-related pathologies^8^, how distinct functional roles of CRTCs contribute to the coordination of organismal longevity remains unclear.

CRTCs lack DNA-binding activity and depend on transcription factor partners to stimulate gene transcription. CRTCs contain a conserved N-terminal coiled-coil domain, the CREB binding domain, required to bind bZIP transcription factors^9^. *C. elegans* possess a single, highly conserved CRTC family member, CRTC-1, which plays a critical role in the modulation of longevity during low energy conditions. When AMP-activated protein kinase (AMPK) is constitutively active, it inactivates CRTC-1 directly by phosphorylation, and this inhibition is a critical step for AMPK-mediated longevity in *C. elegans*. In addition, CRTC-1 acts through its canonical transcriptional partner CREB in AMPK-mediated longevity^2,3^. Here we identify a new role for CRTC-1 in the epigenetic regulation of longevity that is functionally distinct from the mechanism by which this cofactor modulates the effect of AMPK. Furthermore, we demonstrate that CRTC-1 promotes longevity through a single defined domain by coupling two modes of action: transcriptional activation and histone acetylation.

### CRTC-1-binding proteins

To identify CRTC-1 protein interactions, we tagged endogenous CRTC-1 via clustered regularly interspaced short palindromic repeats-associated protein 9 (CRISPR-Cas9) editing to generate a *crtc-1::3xFLAG* strain and performed immunoprecipitation (IP) followed by liquid chromatography coupled to mass spectrometry (LC-MS). We found 137 potential CRTC-1 interacting proteins (**Fig. 1a and Supplementary Table 1**), including known mammalian CRTCs interactors such as the COPII vesicle coat protein required for vesicle formation in ER to Golgi transport the secretory protein SEC-31/SEC31, the serine/threonine protein phosphatase TAX-6/calcineurin, and to a lesser extent, AAK-2/AMPK alpha 2^2,6,10^ (**Fig. 1a**). STRING *in silico* analysis^11^ revealed that CRTC-1 binds proteins with WD40 repeats, including the WD40-repeat protein WDR-5.1/WDR5 (**Fig. 1b**). WDR-5.1, along with the methyltransferases SET-2/SET1 and ASH-2/ASH2L (components of the complex proteins associated with Set1, COMPASS), catalyses trimethylation of lysine 4 on histone H3 (H3K4me3) and regulates lifespan in *C. elegans*^12,13^. H3K4me3-deficient *C. elegans* have activated sterol regulatory-element binding protein (SREBP1)/SBP-1, which in turn drives a transcriptional network that promotes fat accumulation with specific enrichment of monounsaturated fatty acids (MUFAs), which are necessary for lifespan extension^13^. Since CRTC-1 binds WDR-5.1 and we previously showed that CRTC-1 modulates metabolism during AMPK-dependent longevity^2,3^, we reasoned that CRTC-1 could play a novel role in COMPASSdependent longevity.

**Fig. 1.**
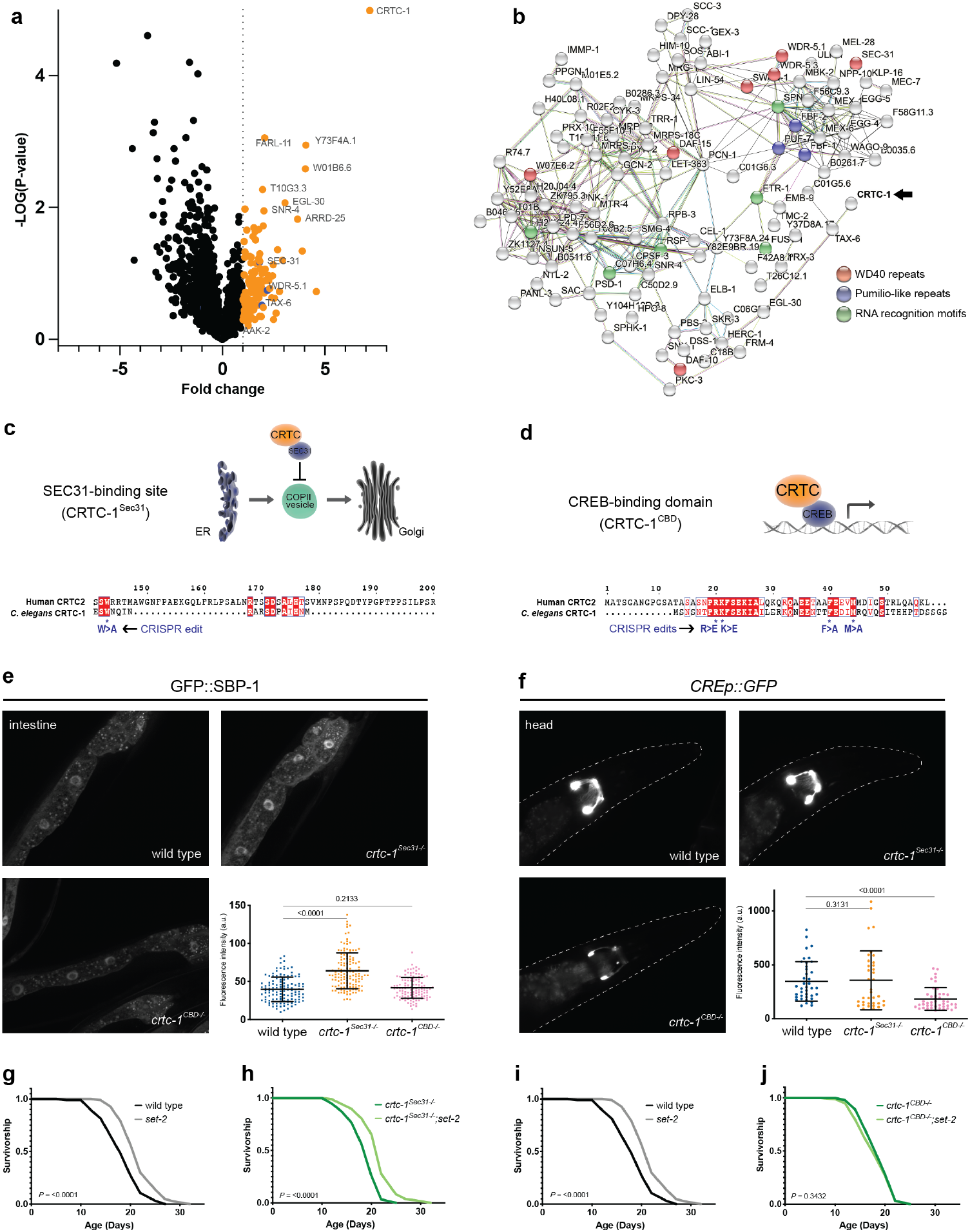
The co-transcriptional function of CRTC-1 is required for H3K4me3-mediated longevity. **a**, Volcano plot showing proteins identified during immunoprecipitation analysis of endogenous CRTC-1::3xFLAG tagged strain relative to wild-type non-tagged strain (N2). The dotted line indicates a one-fold change. Four biological replicates. *P* values by two-sample t-test. See Supplementary Table 1 for IP statistics. **b**, STRING protein-protein interaction network of CRTC-1-binding proteins identified during immunoprecipitation analysis. The network nodes are proteins. The edges represent the predicted functional associations. Redline (presence of fusion evidence), green line (neighbourhood evidence), blue line (co-occurrence evidence), purple line (experimental evidence), yellow line (text mining evidence), light blue line (database evidence), and blackline (co-expression evidence). **c**, Schematic of endoplasmic reticulum (ER) Golgi traffic regulated by the SEC31-binding site of CRTC (top). Amino acid sequence localisation of the SEC31-binding site and CRISPR edit (bottom). **d**, Schematic of co-transcriptional regulation by the CREB-binding domain of CRTC (top). Amino acid sequence localisation of the CREB-binding domain site and CRISPR edits (bottom). **e**, *crtc-1^Sec31-/-^* specifically regulates SREBP1/SBP-1 activation. Intestinal images of GFP::SBP-1 reporter in wild type, *crtc-1^Sec3B-/-^*, and *crtc-1^CBD-/-^* animals. Quantification of GFP::SBP-1 nuclear accumulation; mean ± SEM of n = 123-133 nuclei, pooled from at least three independent experiments, \*\*\*\**P*<0.0001 by Mann-Whitney test. **f**, *crtc-1^CBD-/-^* specifically regulates CRE promoter transcriptional activation. Images of CREp::GFP reporter in neurons of the head in wild type, *crtc-1^Sec31-/-^*, and *crtc-1^CBD-/-^* animals. Quantification of CREp::GFP expression; mean ± SEM of n = 36-44 heads, pooled from at least three independent experiments, \*\*\*\**P*<0.0001 by Mann-Whitney test. **g-h**, Survival curves showing that *set-2(ok952)* mutants live longer than wild type (**g**), and *crtc-1^Sec31-/-^* does not suppress this longevity phenotype (**h**). **i-j**, Survival curves demonstrating that *set-2(ok952)* mutants live longer than wild type (**i**), and *crtc-1^CBD-/-^*completely suppresses this longevity phenotype (**j**). Survival curves compared by the Log-rank (Mantel-Cox) method (see Supplementary Table 2 for lifespan statistics).

### CRTC-1 specifically regulates lifespan extension and MUFA accumulation under H3K4me3 deficiency

We generated a novel *C. elegans crtc-1* null mutant strain (*crtc-1^null^*) to test its role in COMPASS-dependent longevity. *crtc-1^null^* mutants do not show changes in lifespan compared to wild-type animals (**Extended Data Fig. 1a**). As previously reported^12^, RNA interference (RNAi) of H3K4me3 modifiers *set-2, ash-2*, and *wdr-5.1* extends lifespan in wild-type worms (**Extended Data Fig. 1b, d, and f**). In *crtc-1^null^* mutants, *wdr-5.1 RNAi* still increases lifespan by 20% (**Extended Data Fig. 1c**). However, lifespan extension by *ash-2 RNAi* is significantly reduced in *crtc-1^null^* mutants (from 35% to 19%), and the *crtc-1^null^* entirely suppresses *set-2 RNAi* longevity (**Extended Data Fig. 1d-g**). To further explore the role of CRTC-1 in SET-2-dependent longevity, we crossed the *crtc-1^null^* mutant into the long-lived *set-2(ok952)* mutant strain^12,14^. As with *set-2 RNAi,* the absence of CRTC-1 fully suppresses lifespan extension in *set-2* mutants (**Extended Data Fig. 1h-i**).

We next examined whether CRTC-1 is a more general modulator of other longevity pathways. We combined *crtc-1^null^* animals with *raga-1(ok386)* (mTORC1 pathway) and *clk-1(qm30)* (electron transport chain) mutants, reduced insulin/insulin-like growth factor-1 signalling via *daf-2 RNAi,*and dietary restriction^2,3,15–18^. In all conditions, there is a significant increase of lifespan in both wild-type (31%, 40%, 257%, and 39%, respectively) and *crtc-1^null^* (26%, 55%, 263%, and 42%, respectively) backgrounds (**Extended Data Fig. 1j-q**). Therefore, CRTC-1 is not a global longevity regulator but instead is specifically required for H3K4me3-dependent longevity. In addition, since activation rather than deletion of CRTC-1 suppresses AMPK-mediated longevity^2,3^, these data highlight a complex and contextual role for how CRTC-1 activity state can impact healthy ageing.

H3K4me3-deficient *C. elegans* accumulate MUFAs, which is necessary for lifespan extension^13^. We therefore examined whether CRTC-1 regulates MUFAs accumulation in *set-2* mutants by performing gas chromatography coupled to mass spectrometry (GC-MS). As previously reported^13^, there is a significant accumulation of cis-vaccenic acid in *set-2* mutants, and this is decreased in *crtc-1^null^;set-2* double mutants (**Extended Data Fig. 2a**), suggesting that CRTC-1 is required for the accumulation of this MUFA specifically in *set-2* animals.

### Targeting CRTC-1 functions in longevity

CRTCs have multiple functions related to different cellular processes and age-related diseases^8^, and we hypothesised that a specific CRTC-1 functional domain might promote longevity under conditions of H3K4me3 deficiency. We used CRISPR-Cas9 gene editing to selectively inhibit two functions of endogenous CRTC-1 in *C. elegans*: its role in COPII trafficking and as a transcriptional coactivator. Mammalian CRTC2 regulates SREBP1 activation and integrates it into TORC1 activity through COPII vesicle formation and competitive sequestration of Sec31A^6^. In *C. elegans,* SBP-1/SREBP1 activation is required for MUFAs synthesis and longevity in H3K4me3-deficient animals^13^. Supporting these data, our immunoprecipitation experiments confirm CRTC-1 interacts with the *C. elegans* counterpart of Sec31A, SEC-31 (**Fig. 1a-b**). Using CRISPR-Cas9, we specifically mutated the amino acid in CRTC-1 required for Sec31A-binding^6^ (**Fig. 1c**). We previously showed that a constitutively activated nuclear form of CRTC-1 suppresses AMPK-dependent longevity by promoting CRH-1/CREB-dependent transcription^2,3,19^, highlighting the transcriptional role of CRTC-1 in longevity. Therefore, separately, we inhibited CRTC-1’s transcriptional coactivator activity by generating point mutations in the CREB binding domain (CBD) (**Fig 1d**).

We used *in vivo* reporters for SBP-1 processing and transcription activation^2,20^ to confirm that the Sec31 and CBD (**Fig. 1c-d**) edits indeed uncouple specific CRTC-1 functions. Using a GFP::SBP-1 reporter, we quantified SBP-1 nuclear accumulation in intestinal cells as an indication of its activation^13^. As expected, the *crtc-1^Sec31-/-^* mutant increases SBP-1 activation (**Fig. 1e**) due to disruption of CRTC-1:Sec31 interaction. To assess CRTC-1 transcriptional coactivator activity, we used a CREB transcriptional reporter, a cAMP response element sequence fused with GFP (CREp::GFP)^2^. The CREp::GFP reporter is mainly expressed in head neurons, and its expression is specifically reduced in the *crtc-1^CBD-/-^* mutant (**Fig. 1f**). Critically, these two CRTC-1 functions can be specifically uncoupled since *crtc-1^CBD-/-^* mutants do not increase SBP-1 activation, and *crtc-1^Sec31-/-^* mutants do not reduce CREp::GFP transcriptional reporter expression (**Fig. 1e-f**). Our data confirm that the roles of CRTC-1 in transcriptional regulation and ER-to-Golgi trafficking are conserved in *C. elegans* and, importantly, that _s_ we can selectively and specifically uncouple these functions in endogenous CRTC-1.

To define the specific CRTC-1 function that regulates H3K4me3-dependent longevity, we crossed the *crtc-1^CBD-/-^* and *crtc-1^Sec31-/-^* strains into *set-2* long-lived mutants and analysed their role in lifespan. Strikingly, although H3K4me3-deficient animals promote MUFAs accumulation through SBP-1 activation^13^, the Sec31 point mutation does not suppress longevity in these animals (**Fig. 1g-h**). In contrast, the CBD point mutation fully suppresses lifespan extension in *set-2* worms (**Fig. 1i-j**). Additionally, *crtc-1^CBD-/-^* is required to promote cis-vaccenic MUFA accumulation (**Fig. 2a**). Collectively, these data indicate the co-transcriptional role of CRTC-1 through its CBD domain (hereafter named CRTC-1^CBD^, referring to the functional wild-type coactivator domain) regulates - H3K4me3-dependent longevity. Contrary to AMPK longevity, where the CRTC-1-mediated transcriptional activation suppresses longevity^2,3^, H3K4me3 deficiency (where overall transcription levels are decreased) requires transcriptional activation driven by CRTC-1^CBD^ to extend lifespan. Therefore, our data suggest that transcriptional activation is necessary to promote longevity in H3K4me3-deficient animals.

**Fig. 2.**
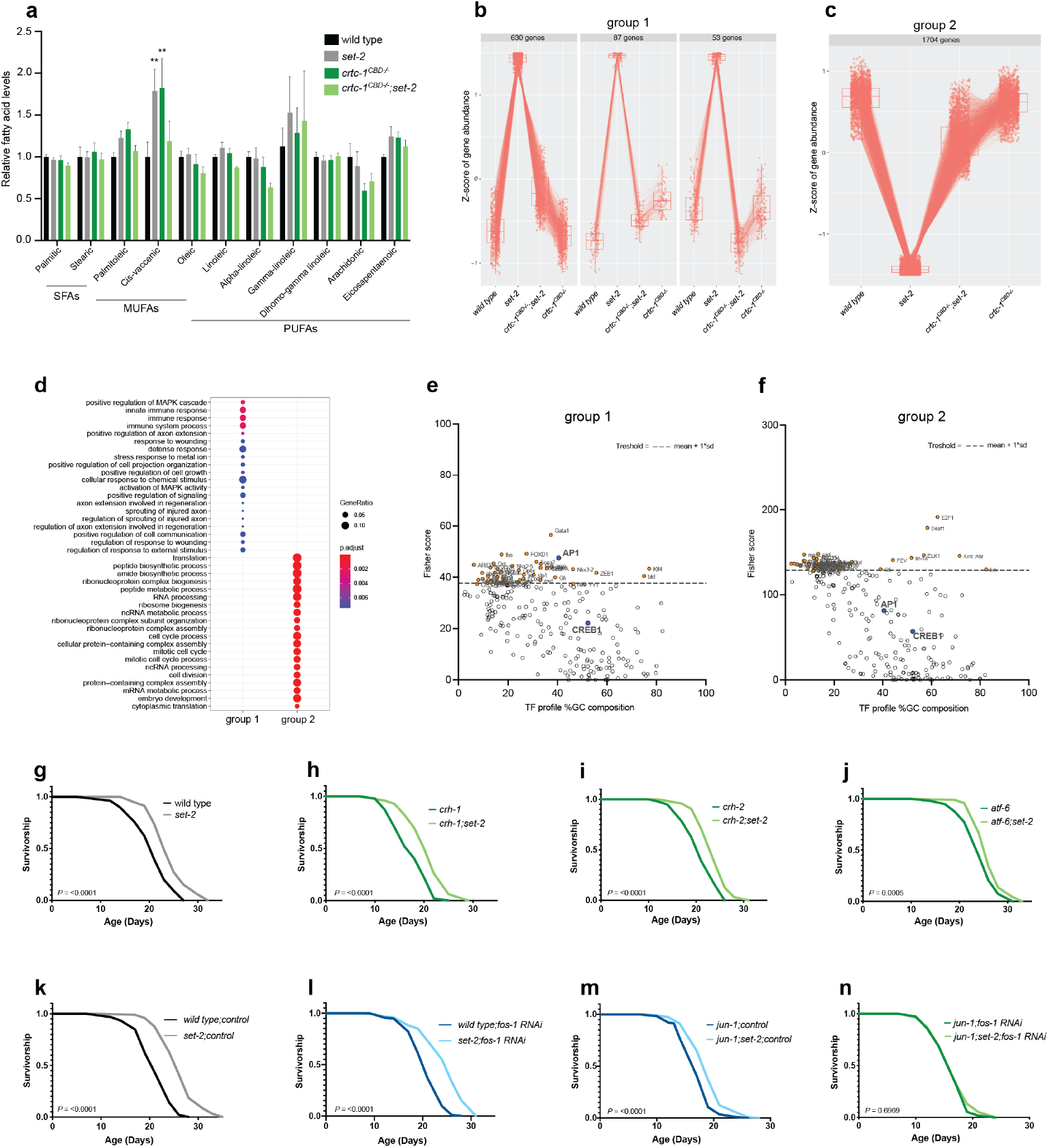
CRTC-1^CBD^ mediates specific longevity pathways and acts through AP-1 to extend lifespan in H3K4me3-deficient animals. **a**, GC-MS quantification of fatty acids. Mean ± SEM, \*\**P*<0.01 by unpaired two-way ANOVA. **b-c**, Cluster analysis identifies 770 genes that show increased expression in *set-2(ok952)*, which is reversed in the double *crtc-1^CBD-/-^::set-2(ok952)* mutant (**a**) and 1704 genes that show decreased expression in *set-2(ok952)* and reversed in the absence of CRTC-1^CBD^ (**b**). **d**, Over-representation analysis of the Gene Ontology biological process terms for genes comprising each cluster shown in panels (**a-b**). **e-f**, Single site analyses to detect over-represented conserved transcription factor binding sites in groups 1 (**e**) and 2 (**f**). Analysis by oPOSSUM version 3.0. **g-j**, Survival curves showing that the lifespan extension in *set-2(ok952)* (**g**) mutants does not require the transcriptional factors *crh-1(n3315)* (**h**), *crh-2(gk3293)* (**i**), and *atf-6(ok551)* (**j**). **k-n**, AP-1 components, *fos-1* and *jun-1,* do not suppress *set-2(ok952)* longevity separately (**k-m**), but the absence of the entire complex (*jun-1(gk557);fos-1RNAÏ)* fully suppresses *set-2(ok952)* longevity (**n**). Survival curves compared by the Log-rank (Mantel-Cox) method (see Supplementary Table 2 for lifespan statistics).

### CRTC-1^CBD^ regulates specific longevity-related pathways

Our data indicate that the co-transcriptional role of CRTC-1 is necessary to extend lifespan in H3K4me3-deficient animals (**Fig. 1i-j**). Next, we reasoned that CRTC-1^CBD^-dependent transcriptional changes in *set-2* mutants might reveal mechanistic insight into how CRTC-1 regulates COMPASS-mediated longevity. Via RNA sequencing, we identified CRTC-1^CBD^-dependent differentially expressed genes (DEGs) in *set-2* animals that cluster into two main groups: genes upregulated in *set-2* and downregulated by *crtc-1^CBD-/-^* in the double mutant (group 1) and vice versa, genes downregulated in *set-2* mutants and then upregulated by addition of *crtc-1^CBD-/-^* (group 2) (**Fig. 2b-c and Supplementary Table 3**). Functional analysis within these two clusters reveals that Gene Ontology (GO) terms between the groups are distinct (**Fig. 2d, Extended Data Fig. 2b-c, and Supplementary Table 3**). Group 1 DEGs are enriched for processes involved in activation of the MAPK cascade, including immune and stress responses, whereas group 2 includes GO terms related to translation, including ribosome biogenesis, ribonucleoprotein assembly, and RNA processing (**Fig. 2d**). Importantly, both activation of MAPK cascade and reduction of translation are defined signatures of longevity^21–23^, and CRTC-1^CBD^ specifically regulates expression of genes involved in these processes. CRTC-1^CBD^ does not regulate transcriptional changes directly linked to lipid metabolism in either group 1 or 2 (**Fig. 2d**). Thus, even though CRTC-1^CBD^ alters the composition of some MUFAs, this effect does not seem to be via direct transcriptional regulation of genes that control fatty acid synthesis. Together, our data suggest that instead of a generalised dysregulation of transcription by the absence of CRTC-1^CBD^ in H3K4me3-deficient animals, CRTC-1 ^CBD^-dependent transcriptional regulation of genes in specific pathways is coupled to longevity.

### AP-1 regulates longevity in H3K4me3-deficient animals

The CBD domain of CRTC-1 is critical to bind multiple bZIP factors^19^, including CREB, the heterodimer activator protein 1 (AP-1), and the activating transcription factor (ATF)6^9,24–26^. Using our transcriptional data (**Fig. 2b-c**), we performed Single Site Analysis to detect over-represented conserved transcription factor binding sites^27–29^ in our DEG groups. CREB1 and AP-1 transcription factors have gene targets in both groups (**Fig. 2e-f and Supplementary Table 4**). However, only AP-1 is above the significance threshold in group 1, meaning DEGs upregulated in H3K4me3-deficient animals and downregulated by the absence of CRTC-1^CBD^ (**Fig. 2b**) are significantly enriched for direct targets of AP-1 (**Fig. 2e and Supplementary Table 4**). To identify CRTC-1 binding proteins that are specifically dependent on its CBD domain, we compared peptides bound to the endogenous CRTC-1::3xFLAG versus endogenous CRTC-1^CBD-/-^::3xFLAG strains in the *set-2* mutant background by LC-MS (**Extended Data Fig. 3a-b and Supplementary Table 5**). After removing background, 1062 peptides bound to CRTC-1::3xFLAG and 1127 bound to CRTC-1 ^CBD-/-^::3xFLAG, of which 896 are shared. We focused on peptides specifically bound to CRTC-1 when the CBD is functional. Previously described transcription factors are not identified in the analysis, perhaps due to transient interactions, but interestingly the CRTC-1^CBD^ binds mainly to chromatin proteins in *set-2* mutants (**Extended Data Fig. 3b**). This supports our model where CRTC-1 is required in the nucleus to modulate H3K4me3 longevity.

To test the causal role of CRTC-1 transcription factor partners in H3K4me3-dependent longevity, we used a combination of mutants and RNAi for their worm counterparts: CRH-1/CREB1, CRH-2/CREB3L1, ATF-6/ATF6, and FOS-1: JUN-1/AP-1^9,24–26^. Although CRTC-1 governs AMPK-mediated longevity and metabolism via the CREB transcription factor^3^, CRH-1 and CRH-2 are dispensable for longevity in *set-2* mutants (**Fig. 2g-i**). The *atf-6(ok551)* mutant by itself is long-lived^30^. *set-2* deletion significantly increases lifespan in *atf-6* mutants (8%), although to a lesser extent than in wild type (14%) (**Fig. 2j**). Separately, inactivation of AP-1 components, JUN-1 and FOS-1 alone do not completely suppress *set-2* longevity (**Fig. 2k-m**), likely due to redundancy with other AP-1 components^31^. Strikingly, however, co-inhibition of both JUN-1 and FOS-1 together (*jun-1(gk557);fos-1 RNAi)* fully suppresses *set-2* longevity (**Fig. 2n**). These data indicate that the complete heterodimer AP-1 transcription factor is necessary to extend longevity and point to the CRTC-1^CBD^:AP-1 complex as the critical transcriptional core that promotes longevity in H3K4me3-deficient animals.

### CRTC-1^CBD^ regulates lifespan cell nonautonomously

H3K4me3 modifiers function in the germline to promote longevity^12,13^, and CRTC-1 is known to mediate longevity from neurons during AMPK activation^3^. Our data here show that CRTC-1^CBD^ is required to extend lifespan of *C. elegans* with an H3K4me3 deficiency (**Fig. 1**). We therefore asked whether inter-tissue signalling might be necessary for the role of CRTC-1^CBD^ in H3K4me3 longevity. First, we performed tissue enrichment analysis of the DEGs present in the two CRTC-1^CBD^ dependent groups obtained from our RNA-Seq data (**Fig. 2b-c**). Group 1 has significant enrichment of genes expressed in neurons (**Fig. 3a**). In contrast, DEGs in group 2 are enriched in the gonad (**Fig. 3b**). Additionally, we performed tissue enrichment analysis for AP-1 targets from group 1 (**Fig. 2e**). Similar to CRTC-1, AP-1 direct targets from group 1 are enriched for genes expressed in the nervous system (**Fig. 3c**). To explore how the tissue enrichment analysis might link to CRTC-1 tissue localisation, we characterised endogenous expression of CRTC-1 via GFP knock-in at the endogenous locus. CRTC-1::GFP expression is particularly high in neurons, while also present in muscles and intestine but not in the germline (**Fig. 3d**). Taken together, these data indicate that the co-transcriptional role of CRTC-1 in regulating H3K4me3-dependent longevity (**Fig. 1d, f, i, and j**) correlates with transcriptional activation of neuron-enriched genes in group 1 (**Fig. 2b and 3a**) and suggest that CRTC-1 may modulate H3K4me3-dependent longevity cell nonautonomously from the nervous system.

**Fig. 3.**
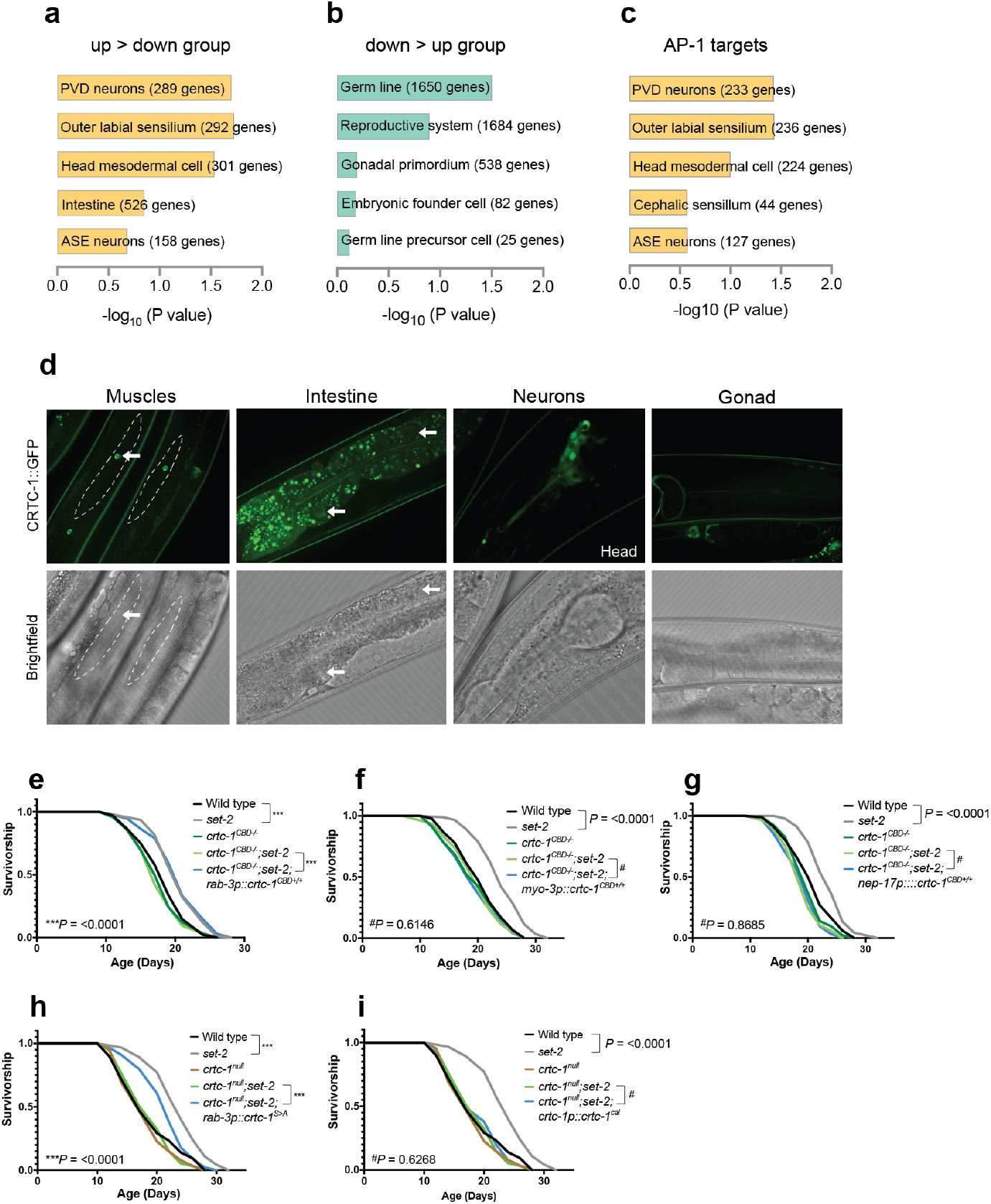
CRTC-1^CBD^ regulates longevity cell nonautonomously. **a-b**, Tissue enrichment analyses for genes comprising groups 1 (**a**) and 2 (**b**). Analysis by WormBase version WS280. **c**, Tissue enrichment analysis of AP-1 target genes from group 1 using WormBase version WS280. **d**, Confocal images showing that endogenous CRTC-1::GFP is expressed in muscles, intestine, and neurons but not in the germline. Arrows point CRTC-1::GFP expression in nuclei from muscles and intestine. **e-g**, Survival analysis demonstrating that lifespan extension in *set-2(ok952)* mutants is suppressed by *crtc-1^CBD-/-^* and significantly rescued by a single copy of neuronal *crtc-1^CBD+/+^* (**e**), but not when *crtc-1^CBD+/+^* is expressed in muscles (**f**) or intestine (**g**). **h-i**, Survival analysis demonstrating that nuclear (**h**), but not cytosolic (**i**), localisation of CRTC-1 is required to extend lifespan in *set-2(ok952)* mutants. Survival curves compared by the Log-rank (Mantel-Cox) method (see Supplementary Table 2 for lifespan statistics).

To directly examine the role of neuronal CRTC-1^CBD^ activity on H3K4me3-dependent longevity, we rescued CRTC-1^CBD^ activity in specific tissues in *crtc-1^CBD^;set-2* mutants. Using our SKI LODGE system^32^, which allows for efficient CRISPR-Cas9-mediated knock-ins regulated by tissue-specific promoters, we inserted a single copy of *crtc-1* cDNA wild-type sequence into an intergenic cassette containing the *rab-3* promoter that drives gene expression specifically in neurons. Expressing CRTC-1 from the *rab-3* promoter fully rescues the lifespan extension in *crtc-1^CBD-/-^;set-2* mutants (**Fig. 3e**). In contrast, expressing CRTC-1 directly in muscles or intestine by single copy transgene, using the muscle-specific *myo-3* or intestine-specific *nep-17* promoters, does not rescue the lifespan extension in the double mutants (**Fig. 3f-g**). We then tested whether nuclear or cytosolic localisation of CRTC-1 is necessary for H3K4me3 longevity. We rescued the *crtc-1^null^* mutant with neuronal expression of a constitutively nuclear form of *crtc-1* (S76A, S179A, referred to as *crtc-1^S>A^*)^3^. Neuronal and nuclear expression of CRTC-1^S>A^ significantly rescues the lifespan extension in *crtc-1^null^;set-2* mutants (30% compared to 40% in single *set-2* mutants versus wild-type animals) (**Fig. 3h**). In contrast, cytosolic restricted CRTC-1, achieved by a mutation in the calcineurin binding site (*crtc-1^cal^*)^3^ which prevents activation by dephosphorylation, does not rescue lifespan extension (**Fig. 3i**). Taken together, CRTC-1^CBD^ nuclear activity specifically in neurons therefore modulates longevity in animals with H3K4me3 deficiency.

### Histone acetylation promotes longevity

Next, we explored how CRTC-1 impacts histone modifications in the context of COMPASS-mediated longevity. Mammalian CRTC2 associates with the lysine acetyltransferase 2B KAT2B (p300/CBP-associated factor PCAF-1 in *C. elegans*), which in turn acetylates lysine 9 on histone H3 (H3K9ac) at promoters of gluconeogenic genes, upregulating transcription^4^. CRTC2 also interacts with the acetyltransferases adenovirus E1A-associated 300-kD protein (p300) and CREB-binding protein (CBP)^5^. The *C. elegans* ortholog of the mammalian p300 and CBP, the putative histone acetyltransferase CBP-1, was recently shown to affect H3K18ac and H3K27c levels^33^. Therefore, we sought to explore whether CRTC-1^CBD^ mediates H3K9ac, H3K18ac, and H3K27ac marks in *set-2* mutants. By Western blotting with histone mark-specific antibodies, all three acetylation marks are slightly increased in *set-2* mutants: H3K27ac (−5%), H3K18ac (+25%), and H3K9ac (+11%). The same histone marks are then reduced in single *crtc-1^CBD-/-^* mutants: H3K27ac (-5%), H3K18ac (−14%), and H3K9ac (−16%). Strikingly, however, acetylation is dramatically reduced in the double *crtc-1^CBD-/-^;set-2* mutants: H3K27ac (−45%), H3K18ac (−34%), and H3K9ac (−32%) (**Fig. 4a and Extended Data Fig. 4b-d**). In contrast, *crtc-1^CBD-/-^* does not change H3K4me3 levels in *set-2* mutants (**Fig. 4a and Extended Data Fig. 4a**). These results indicate that the CRTC-1^CBD^ regulates H3K9ac, H3K18ac, and H3K27ac marks, particularly in H3K4me3-deficient animals.

**Fig. 4.**
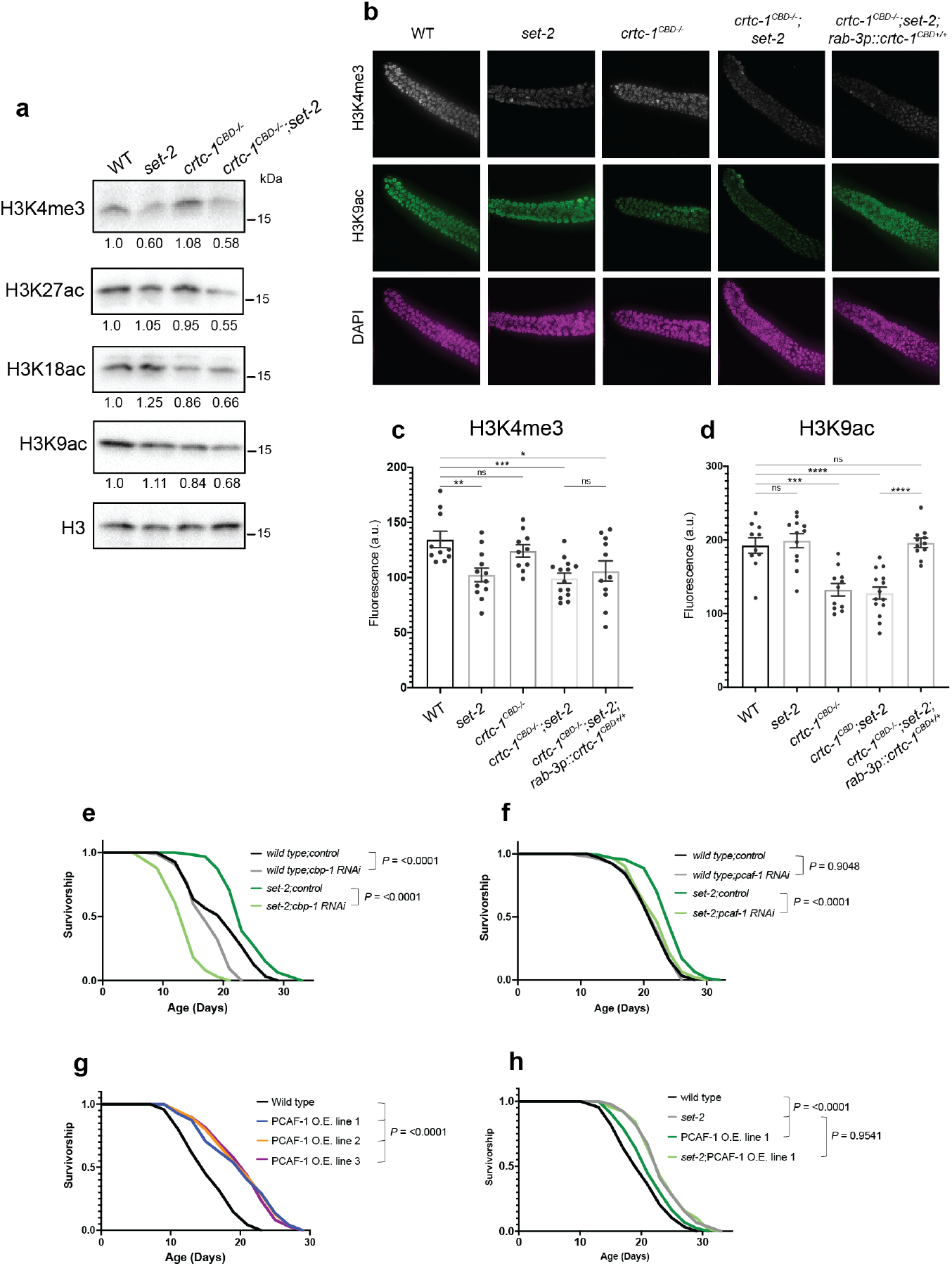
CRTC-1^CBD^ maintains histone acetylation levels, and these are necessary for lifespan extension. **a**, H3K9ac, H3K18ac, and H3K27ac marks are reduced in a CRTC-1^CBD^-dependent manner under H3K4me3 deficiency induced by a loss of function of *set-2*. Western blots of wild-type, *set-2(ok952), crtc-1^CBD-/-^*, and *crtc-1^CBD-/-^;set-2(ok952)* worms. H3 used as loading control and values represent the mean of three independent blots. See Extended Data Fig. 4 for Western blot quantifications. **b**, Fluorescence images of the distal region of *C. elegans* gonad from wild-type, *set-2(ok952), crtc-1^CBD-/-^, crtc-1^CBD-/-^;set-2(ok952)* and *crtc-1^CBD-/-^;set-2(ok952);rab-3p::crtc-1^CBD+/+^* worms co-immunostained for H3K4me3 (grey), H3K9ac (green), and DAPI (magenta). **c**, Quantification of fluorescence intensity detected for H3K4me3 (**a**). *set-2* and *crtc-1^CBD-/-^;set-2* worms show a significant reduction of the H3K4me3 mark, which is not rescued in the *crtc-1^CBD-/-^;set-2(ok952);rab-3p::crtc-1^CBD+/+^* strain. Mean ± SEM of n = 10-14 gonads, pooled from three independent experiments, \**P*<0.05, \*\**P*<0.01, and \*\*\**P*<0.001 by unpaired t-test. ns = not significant. **d**, Quantification of fluorescence intensity detected for H3K9ac (**a**). *crtc-1^CBD-/-^* and *crtc-1^CBD-/-^;set-2(ok952)* worms show a significant reduction of the H3K9ac mark, which is rescued in the *crtc-1^CBD-/-^;set-2(ok952);rab-3p::crtc-1^CBD+/+^* strain. Mean ± SEM of n = 10-14 gonads, pooled from three independent experiments, \*\*\**P*<0.001 and \*\*\*\**P*<0.0001 by unpaired t-test. ns = not significant. **e-f**, Survival curves demonstrating that lifespan extension in *set-2(ok952)* mutants is suppressed by *cbp-1* (**e**) and *pcaf-1* (**f**) RNAi. (**g-h**) Survival curves demonstrating that overexpression (OE) of PCAF-1 increases wild-type lifespan in three different lines (**g**) and does not further extend the lifespan in *set-2(ok952)* mutants (**h**). Survival curves compared by the Log-rank (Mantel-Cox) method (see Supplementary Table 2 for lifespan statistics).

Given that H3K4me3 modifiers promote longevity from the germline^12,13^ and CRTC-1^CBD^ modulates H3K4me3 longevity from neurons (**Fig. 3**), we asked whether CRTC-1^CBD^ could regulate histone acetylation cell nonautonomously. Supporting our Western blot analyses (**Fig. 4a**), immunofluorescence confirms that *crtc-1^CBD-/-^* does not change H3K4me3 levels in the gonad (**Fig. 4b-c**), but H3K9ac levels are significantly reduced in both *crtc-1^CBD-/-^* and *crtc-1^CBD-/-^;set-2* animals (**Fig. 4b and d**). Interestingly, restoring *crtc-1^CBD+/+^* expression only in neurons fully rescues H3K9ac germline levels in *crtc-1^CBD-/-^;set-2* mutants (**Fig. 4b and d**) but does not affect reduced levels of H3K4me3 in the gonad (**Fig. 4b-c**). Therefore, similarly to lifespan, CRTC-1^CBD^ regulates H3K9ac cell nonautonomously from neurons. We then tested the role of AP-1 on histone acetylation. As with the *crtc-1^CBD-/-^* results, inactivation of AP-1 induces a significant reduction of the H3K9ac mark in the gonad but an increase in global levels (**Extended Data Fig. 4e-i**), indicating a dual role of AP-1 on histone acetylation in soma and germline.

So far, our data indicate that CRTC-1^CBD^ regulates both lifespan and histone acetylation from neurons. Therefore, we sought to further define the potential neuronal pathways by which CRTC-1 might modulate both longevity and histone acetylation. Tissue-enrichment analysis reveals that PVD mechanosensory neurons are the most significant tissue in group 1 DEGs (**Fig. 3a**). We also saw transcriptional changes in components of the acetylcholine pathway (**Supplementary Table 3**), which controls response to mechanical stimuli in PVD neurons^34^. As expected, mutations in core components of the acetylcholine pathway are lethal^35^. To test causal roles of this pathway in longevity and histone modification, we selected mutants to genes that show mild defects, including members of vesicle formation (*sup-1*) and acetylcholinesterases (*ace-1, ace-2*, and *ace-3*)^35,36^. Mutants in these components have a global and germline increase in H3K9ac and exhibit significant lifespan extension, ranging from 23% to 42% (**Extended Data Fig. 5a-i**). Moreover, *set-2* knockdown does not further increase lifespan in any of these mutants (**Extended Data Fig. 5f-h**). It is also possible to alter histone acetylation by disrupting the production of acetyl-CoA, the primary source of histone acetylation^37^. Acetate can enter the cell by monocarboxylic acid transporters and subsequently be converted to acetyl-CoA in the mitochondria^38^. Knockdown of a putative monocarboxylic acid transporter in *C. elegans*, *mct-6*, reduces global and germline H3K9ac levels, does not increase lifespan alone and does not further extend lifespan in *set-2 RNAi* animals (**Extended Data Fig. 5a-e and i**). Taken together, our data suggest that CRTC-1^CBD^ regulates histone acetylation through cholinergic signals, notably when H3K4me3 levels are reduced, and that histone acetylation contributes to lifespan extension in H3K4me3-deficient animals.

To test the hypothesis that histone acetylation is required for longevity during H3K4me3 deficiency, we knocked down the *C. elegans* orthologs of the histone acetyltransferases associated with CRTCs in mammals that also regulate the acetylation marks mentioned above, PCAF-1/KAT2B and CBP-1/p300/CBP^4,5^. Supporting our previous findings, RNAi knockdown of the putative histone acetyltransferases *cbp-1* and *pcaf-1* entirely suppresses longevity in *set-2* mutants (**Fig. 4e-f**). Remarkably, *cbp-1* knockdown has a dramatic effect on *set-2* mutants. The lifespan in *set-2;cbp-1 RNAi* is shorter (−40%) than *wild type;cbp-1 RNAi* lifespan (−23%) (**Fig. 4e**), indicating that CBP-1 plays a more essential role in H3K4me3-deficient animals than animals with wild-type H3K4me3 levels.

Our data demonstrate that histone acetylation is required to extend lifespan in the context of H3K4me3 deficiency. Finally, we reasoned that if histone acetylation positively affects ageing, lifespan would be increased by overexpressing histone acetyltransferases associated with CRTCs. Indeed, overexpression of PCAF-1 is sufficient to extend lifespan in a wild-type background by up to 40% (**Fig. 4g**) and does not further extend lifespan in *set-2* mutants (**Fig. 4h**). Collectively, these data suggest that CRTC-1^CBD^ mediates longevity during H3K4me3 deficiency via remodelling histone acetylation levels, and remarkably, that this acetylation is sufficient to promote longevity in wild-type worms, identifying this histone modification as a novel marker for longevity.

## DISCUSSION

CRTCs have several functions and have been implicated in many age-related diseases such as cancer, neurodegeneration, diabetes, and other metabolic disorders^8,39^. Thus, CRTCs are key integrators that regulate cellular signalling, and their functionality is context-dependent. Linking CRTC functions to specific physiological processes will, therefore, offer novel opportunities for clinical therapies. We previously demonstrated that an overexpressed active form of CRTC-1 in neurons suppresses lifespan extension when AMPK is constitutively activated in the entire worm^3^. Conversely, in this study, we found a positive role for CRTC-1 in regulating longevity due to H3K4me3 deficiency. While CRTC-1 modulates genes associated with metabolism and mitochondria under AMPK activation to suppress longevity^3^, we found that CRTC-1^CBD^ regulates genes involved in immune and stress responses in H3K4me3-deficient animals to extend lifespan. Other positive and negative roles have been described for CRTCs in cancer, diabetes, and autoimmune disease^8,39^. Here, we discover a dual role of CRTC-1 in ageing, supporting the idea that CRTC’s roles in maintaining homeostasis are context-dependent.

We propose a model in which lifespan extension induced by H3K4me3 methyltransferase deficiency requires a specific function (co-transcriptional regulation) of CRTC-1 driven by its CBD domain, which in turn triggers a neuron-to-periphery cholinergic signal via the transcription factor AP-1, leading to activation of gene expression and maintenance of histone acetylation levels. Surprisingly, although H3K4me3 deficiency couples lipid metabolism to longevity through the SBP-1/SREBP1 transcription factor in worms^13^ and mammalian CRTC2 regulates lipid metabolism by inducing COPII trafficking-dependent activation of SREBP1^6^, we demonstrate that this CRTC-1 function is not required for H3K4me3 longevity. Although we have demonstrated that a deficiency in H3K4me3 promotes longevity^12,13,40^, recent work suggests that this deficiency shortens lifespan^41^. However, the food source differs from our conditions, and in addition, RNAi in *set-2* extends lifespan in wild-type worms^12–14,40^.

How do global changes in histone acetylation regulate ageing? Historically, one of the most studied families of deacetylases are the sirtuins. Extensive work suggests that sirtuins suppress age-related pathologies and promote healthspan^42^. Increased expression levels of sirtuin deacetylases extend lifespan of *S. cerevisiae, C. elegans, D. melanogaster*, and mice^43–46^, indicating that reduction of acetylation has a positive effect on aging. However, in recent years, several studies have demonstrated that increased acetylation also promotes longevity. The acetyltransferase CBP-1/p300/CBP and the MYST family histone acetyltransferase (MYS-1/TRR-1) promote longevity in worms, regulating mitochondrial stress response and upregulating the DAF-16/FOXO transcription factor, respectively^33,47^. In addition, silencing the deacetyltransferase HDA-6/HDAC6 increases lifespan in worms and flies^48^. Further, depletion of CBP-1, MYS-1/TRR-1, and HDA-6 is associated with immune response and stress resistance^33,47,48^. We found that these pathways are specifically activated in a CRTC-1^CBD^-dependent manner, suggesting conserved mechanisms in long-lived animals with active histone acetylation.

Although we found that CRTC-1 binds WDR-5.1, surprisingly, CRTC-1 is not required for WDR-5.1-mediated longevity. Beyond histone trimethylation, WDR-5.1/WDR5 also associates with histone acetyltransferase complexes in mammals and worms^49–51^. Further, it has been shown that WDR-5.1/WDR5 modulates other histone marks besides H3K4me3, including H3K9me2 and H3K9ac^4,52^. Thus, WDR-5.1 likely regulates longevity by affecting H3K4me3 levels and regulating histone acetylation. This theory may explain why CRTC-1 is not required for WDR-5.1-mediated longevity.

It remains unclear how CRTC-1^CBD^ regulates histone acetylation. One of the primary sources of an acetyl group for histones is the metabolite acetyl-CoA, and it has been shown that cellular metabolism is intertwined with chromatin dynamics since many metabolites constitute histone and DNA modifications^53^. In addition, the acetylcholine pathway produces acetyl-CoA as an intermediary metabolite, and we show here that this pathway also modulates histone acetylation and lifespan. Since CRTCs have been associated with metabolic and neuronal pathways^8^, it is possible that CRTC-1^CBD^ regulates histone acetylation by controlling metabolic intermediaries that act cell nonautonomously. Beyond global changes in histone acetylation or trimethylation, which genes are targeted by these marks in specific contexts and tissues in order to promote longevity remains unknown. Therefore, it will be critical to identify components that integrate histone marks with gene expression, such as CRTC-1. Finally, if promoting histone acetylation induces longevity, as our data indicate, selectively targeting histone acetylation modifiers in specific tissues may be sufficient to generate an organismal response that promotes healthier ageing.

## ACKNOWLEDGMENTS

We thank the Caenorhabditis Genetics Center for providing worm strains. We thank Zon Weng Lai from the Harvard Chan Advanced Multiomics Platform (ChAMP) for the mass spectrometry analysis for immunoprecipitation. We are also grateful to members of the Mair laboratory for helpful discussions on the project and the manuscript.

## AUTHOR CONTRIBUTIONS

CGSG and WBM designed the study. CGSG performed most experiments and analysed results. LILL performed immunostainings and image analysis in MPC lab. KP performed gas chromatography/mass spectrometry analysis of fatty acid profiles in AB lab. CH, AP, CM, LPT and AS performed lifespan repeats. JL contributed to analysing the RNA-sequencing data. CGSG and WBM wrote the manuscript and implemented comments and edits from all authors.

## METHODS

### Materials & Correspondence

Further information and requests for resources and reagents should be directed to and will be fulfilled by the Lead Contact, William Mair. All unique and stable reagents generated in this study are available from public repositories (e.g., Caenorhabditis Genetics Center for *C. elegans* strains) or from the Lead Contact with a completed Materials Transfer Agreement. Source transcriptomic data will be available, and other original data will be available upon request.

### *C. elegans* strains and husbandry

N2 wild-wild type, MQ130 [*clk-1(qm30)*], VC222 [*raga-1(ok952)*], RB772 [*atf-6(ok551)*], VC3149 *[crh-2(gk3293)*], VC1200 [*jun-1(gk557)*], RM3571 *[sup-1(e995 e2636)*], PR1300 [*ace-3(dc2)*], and GG201 [*ace-2(g72);ace-1(p1000)*] *C. elegans* strains were obtained from the Caenorhabditis Genetic Center, which is funded by NIH Office of Research Infrastructure Programs (P40 OD010440). The *crh-1(n3315)* mutant was provided by Mark Alkema. Worms were maintained on standard nematode growth media (NGM) seeded with *E. coli* OP50-1 and maintained at 20 °C.

### Microbe strains

OP50-1 bacteria were cultured overnight in LB at 37 °C, after which 100 ml of liquid culture was seeded on plates to grow for two days at room temperature. Unless otherwise noted, worms were grown at 20 °C on the strain *E. coli* HT115 (empty vector, EV) for all experiments. RNAi experiments employed HT115 bacteria from the Ahringer library (Source Bioscience) expressing dsRNA against the gene noted or empty vector control. HT115 bacteria were cultured overnight in LB containing 100 mg ml^-1^ carbenicillin and 12.5 ml ml^-1^ tetracycline at 37 C, after which 100 mL of LB was seeded on NGM plates containing 100 mg ml^-1^ carbenicillin to grow for two days at room temperature. dsRNA expression was induced by adding 100 ml IPTG (100 mM) at least 2 hours before worms were introduced to the plates. RNAi experiments for *ash-2*, *set-2*, and *wdr-5.1* were performed as previously described^13^.

### Microinjection and CRISPR-Cas9-triggered homologous recombination

All CRISPR edits and insertions required to generate the strains were performed using the previously described CRISPR protocol^32,54^. Briefly, homology repair templates were amplified by PCR, using primers that introduced a minimum stretch of 35 bp homology at both ends. Single-stranded oligo donors (ssODN) were also used as repair templates. CRISPR injection mix reagents were added in the following order: 0.375 μl Hepes pH 7.4 (200 mM), 0.25 μl KCl (1 M), 2.5 μl tracrRNA (4 μg/μl), 0.6 μl *dpy-10* crRNA (2.6 μg/μl), 0.25 μl *dpy-10* ssODN (500 ng/μl), and PCR or ssODN repair template(s) up to 500 ng/μl final in the mix. Water was added to reach a final volume of 8 μl. 2 μl purified Cas9 (12 μg/μl) added at the end, mixed by pipetting, spun for 2 min at 13000 rpm and incubated at 37 °C for 10 min. Mixes were microinjected into the germline of day 1 adult hermaphrodite worms using standard methods^55^.

### Single-Copy Transgene Construction

*crtc-1* and *jun-1* single-copy tissue-specific rescue strains were generated by CRISPR according to^32^. Specifically, a homology repair template (HR) containing *crtc-1* and *jun-1* cDNA sequence was amplified from plasmid pIM41 and wild-type cDNA, respectively. CRISPR mixes containing the HR templates were prepared according to^32^ and injected into the WBM1141, WBM1126, and WBM1478 strains. The resulting CRISPR-edited alleles were outcrossed 6 times to N2 before use for experiments.

### Survival Analyses

Lifespan experiments were performed as described previously^3,56^. Unless otherwise noted, all experiments were performed on standard NGM containing 100 mg ml^-1^ carbenicillin at 20 °C and on HT115 (empty vector, EV). RNAi experiments were performed from hatching in all cases except for *cbp-1 RNAi*, which started on day 1 of adulthood, on standard NGM containing carbenicillin (100 mg ml^-1^). Expression of dsRNA was induced by pipetting 100 ml IPTG solution (100 mM, containing 100 mg ml^-1^ carbenicillin and 12.5 ml ml^-1^ tetracycline) onto HT115 lawns prior to placing worms. Worms were synchronised by bleaching using gravid adults. After bleaching, embryos were placed on plates. When the progeny reached adulthood (~72 h), 100 worms were transferred to fresh plates with 20 worms per plate. Worms were transferred to fresh plates every other day until reproduction had ceased (day 9-12). Survival was scored every 1–2 days, and a worm was deemed dead when unresponsive to 3 taps on the head and tail. Worms were censored due to contamination on the plate, leaving the NGM, eggs hatching inside the adult, or loss of vulval integrity during reproduction. Lifespan analysis was performed using GraphPad Prism.

### RNA Sequencing

More than 2000 day-one adults were used for each sample. Four biological replicate samples were collected for each genotype. Worms were collected in M9 buffer and washed three times in M9 buffer to remove bacteria. Liquids were removed after last centrifugation, and QIAzol lysis reagent (Qiagen, 79306) was added to each sample before snap-freezing in liquid nitrogen. All samples were stored in a −80 °C freezer until RNA extraction. To break the worm cuticle and improve RNA yield, all samples underwent five freeze-thaw cycles. In each cycle, samples were thawed at 37 °C and then snap-frozen in liquid nitrogen. RNA extraction was performed using QIAGEN RNeasy Mini Kit (QIAGEN, 74104) following manufacturer’s instructions. RNA quality was confirmed using TapeStation system (Agilent Technologies). mRNA libraries were prepared using KAPA mRNA HyperPrep, poly-A selection (Roche) following manufacturer’s instructions. Library quality was checked using TapeStation system (Agilent Technologies). Libraries were then pooled and sequenced with Illumina NovaSeq SP Single Lane using 50 read length, pair-end settings.

### Gene Expression and Functional Analysis

All samples were processed using an RNA-seq pipeline implemented in the bcbio-nextgen project (https://bcbio-nextgen.readthedocs.org/en/latest/). Raw reads were examined for quality issues using FastQC (http://www.bioinformatics.babraham.ac.uk/projects/fastqc/) to ensure library generation and sequencing data were suitable for further analysis. Reads were aligned to the Ensembl94 build of the *C. elegans* genome using STAR^57^. Quality of alignments was assessed by checking for evenness of coverage, rRNA content, genomic context of alignments, complexity, and other quality checks. Using Salmon^58^, expression quantification was performed to identify transcript-level abundance estimates and then collapsed down to the gene level using the R Bioconductor package tximport^59^. Principal components analysis (PCA) and hierarchical clustering methods validated the clustering of samples from the same sample group. Differential expression was performed at the gene level using the R Bioconductor package DESeq2^60^. Differentially expressed genes were identified using the Wald test, and significant genes were obtained using an FDR threshold of 0.05. Significant genes were separated into clusters based on similar expression profiles across the defined sample groups. Gene lists for each cluster were used as input to the R Bioconductor package cluster Profiler^61^ to perform an over-representation analysis of Gene Ontology (GO) biological process terms.

### Microscopy of mounted worms and image analysis

For imaging, one-day-old adult animals were mounted on 2% agarose pads and anaesthetised with 20 mM tetramisole in M9. For visualisation of intestinal nuclear SBP-1 expression, a transgenic line expressing a translational fusion between GFP and SBP-1 was used. For visualisation of CRE reporter expression, a transgenic strain expressing a transcriptional reporter was used, CREp::GFP. In both cases, images were acquired using a Zeiss Axio Imager.M2 microscope equipped with an ApoTome.2 system and an AxioCam MRc camera using identical exposure settings for all experiments. The mean fluorescence intensity in arbitrary units (a.u.) of each intestinal nucleus, for the GFP::SBP-1 reporter, was quantified using the oval brush tool in Fiji ImageJ. The mean fluorescence intensity in arbitrary units of each head, for the CREp::GFP reporter, was quantified using the polygon selection tool in Fiji ImageJ. Then, mean intensity values in arbitrary units were graphed using Graphpad Prism, and statistical significance was determined using a two-tailed Mann-Whitney test. For imaging of endogenous CRTC-1::GFP, images were taken in the Sabri Ulker imaging lab using a Yokogawa CSU-X1 spinning confocal disk system (Andor Technology, South Windsor, CT, USA) combined with a Nikon Ti-E inverted microscope (Nikon Instruments, Melville, NY, USA). Images were taken using a 40× objective lens, Zyla cMOS (Zyla 4.2 Plus USB3) camera, and 488 nm Laser for GFP.

### Immunostainings and image analysis

Whole-mount preparations of dissected gonads and immunostainings were performed as in^62^. Briefly, gonads from 24 h post-L4 hermaphrodites were dissected and fixed with 1% formaldehyde for 5 minutes, freeze-cracked, and post-fixed in ice-cold 100% methanol 1 minute, followed by blocking with 1% BSA for 1 hour. The following primary antibodies were used at the indicated dilutions: mouse α-H3K4me3 (1:500, MAB Institute Inc Wako 305-348191) and rabbit α-H3K9ac (1:1000, Abcam ab10812). The secondary antibodies from Jackson ImmunoResearch Laboratories (West Grove, PA) were used at the following dilutions: α-mouse Cy-3 (1:200) and α-rabbit Alexa 488 (1:500). DAPI was used to counterstain DNA. Vectashield from Vector Laboratories (Burlingame, CA) was used as a mounting media and anti-fading agent. Imaging was performed using an IX-70 microscope (Olympus) with a cooled CCD camera (model CH350; Roper Scientific) controlled by the DeltaVision system (Applied Precision). Images were collected using a 40x objective, and Z-stacks were set at 0.2 μm thickness intervals using identical exposure settings for each antibody. Image deconvolution was done using the SoftWoRX 3.3.6 program (Applied Precision) and processed with Fiji ImageJ. Gonads were imaged by dividing the gonad into three sections from distal to loop regions. To quantify H3K4me3 and H3K9ac expressions, intensity in arbitrary units of each section was quantified using the polygon selection tool in Fiji. Then, all values from each section of a single gonad were pulled together. Mean intensity values in arbitrary units were graphed using Graphpad Prism, and statistical significance was determined by unpaired t-test.

### Co-immunoprecipitation

Pulled-down proteins were obtained from N2 (wild type), *crtc-1::3xFLAG, crtc-1::3xFLAG;set-2(ok952)*, and *crtc-1^CBD-/-^::3xFLAG;set-2(ok952)* immunoprecipitations one-day-old animals. Animals were grown at 20 °C, collected in M9 medium, last two washes were done with lysis buffer with glycerol (100 mM HEPES pH 7.5, 2 mM MgCl_2_, 300 mM, 5 mM 2-Mercaptoethanol, 0.05% NP-40, and 10% glycerol) and protease (Sigma-Aldrich, 8340) and phosphatase (Roche, 04906837001) inhibitors. Liquids were removed after the last centrifugation, and then samples were frozen in liquid nitrogen. Liquid nitrogen-frozen pellets equivalent to 1 g of animals were homogenised in 3 ml of lysis buffer (Sigma-Aldrich, L3412; 50 mM Tris HCl, pH 7.4, 150 mM NaCl, 1 mM EDTA, and 1% TRITON X-100) with protease and phosphatase inhibitors. Worms were lysed via sonication (Qsonica, Q700). Protein concentration was measured using Pierce BCA protein assay kit (Thermo Fisher Scientific, PI23227) following manufacturer’s instructions. Anti-FLAG M2 magnetic beads (Sigma-Aldrich, M8823) were used to immunoprecipitate CRTC-1::3xFLAG. Beads were prepared by washing 3 times with 1x wash buffer (Sigma-Aldrich, W0390; 0.5 M Tris HCl, pH 7.4, and 1.5 M NaCl). Each supernatant from sonication was incubated with pre-washed beads overnight with end-over-end rotation at 4 °C. After incubation, beads were washed 3 times 5 minutes with lysis buffer (Sigma-Aldrich, L3412) on a rotator and protease and phosphatase inhibitors. Next, beads were rewashed 3 times 5 minutes with 1× wash buffer (Sigma-Aldrich, W0390) on a rotator and with protease and phosphatase inhibitors. Then, samples were ready to process for mass spectrometry.

### Mass spectrometry for immunoprecipitation

Pulled-down proteins from *crtc-1::3xFLAG* vs N2 and *crtc-1::3xFLAG;set-2(ok952) vs crtc-1^CBD-/-^::3xFLAG;set-2(ok952)* immunoprecipitations were brought to pH 7.5 with 200 mM HEPES (4-(2-hydroxyethyl)-1-piperazineethanesulfonic acid). Proteins were reduced using 5 mM dithiothreitol (Sigma-Aldrich) at 37 °C for 1 h, followed by alkylation of cysteine residues using 15 mM iodoacetamide (Sigma-Aldrich) in the dark at room temperature for 1 h. Excessive iodoacetamide was quenched using 10 mM dithiotheritol. Protein mixtures were diluted in 1:6 ratio (v/v) using ultrapure water prior to digestion using sequencing grade trypsin (Promega) at 37 °C for 16 h. Subsequently, digested peptides were desalted using self-packed C18 STAGE tips (3 M EmporeTM)^63^ for LC-MS/MS analysis. Desalted peptides were resuspended in 0.1% (v/v) formic acid and loaded onto HPLC-MS/MS system for analysis on an Orbitrap Q-Exactive Exploris 480 (Thermo Fisher Scientific) mass spectrometer coupled to an Easy nanoLC 1000 (Thermo Fisher Scientific) with a flow rate of 250 nl/min. The stationary phase buffer was 0.5% formic acid, and mobile phase buffer was 0.5% (v/v) formic acid in acetonitrile. Chromatography for peptide separation was performed using increasing organic proportion of acetonitrile (5-40% (v/v)) over a 120 min gradient) on a self-packed analytical column using PicoTipTM emitter (New Objective, Woburn, MA) using Reprosil Gold 120 C-18, 1.9 μm particle size resin (Dr Maisch, Ammerbuch-Entringen, Germany). The mass spectrometry analyser operated in data-dependent acquisition mode with a top ten method at a 300-2000 Da mass range.

Mass spectrometry data were processed by MaxQuant software version 1.5.2.8^64^ using the following setting: oxidised methionine residues and protein N-terminal acetylation as variable modification, cysteine carbamidomethylation as fixed modification, first search peptide tolerance 20 ppm, main search peptide tolerance 4.5 ppm. Protease specificity was set to trypsin with up to 2 missed cleavages were allowed. Only peptides longer than five amino acids were analysed, and the minimal ratio count to quantify a protein is 2 (proteome only). The false discovery rate (FDR) was set to 1%for peptide and protein identifications. Database searches were performed using the Andromeda search engine integrated into the MaxQuant environment^65^ against the UniProt *Caenorhabditis elegans* database containing 27,390 entries (November 2020). “Matching between runs” algorithm with a time window of 0.7 min was employed to transfer identifications between samples processed using the same nanospray conditions. Protein tables were filtered to eliminate identifications from the reverse database and common contaminants. Supplementary Tables 1 and 5 show the data used for the Volcano plots (Fig. 1a and Extended Data Fig. 3a) and enriched proteins. Enriched proteins were obtained on basis of their fold change, P value, and presence (log2 intensity values) in each replicate.

### Gas chromatography/mass spectrometry analysis of fatty acid profiles

Quantification of long-chain fatty acids was performed as described^13^. Briefly, for each condition, approximately 500 age-synchronised adults on day 1 were collected in M9 buffer and washed 3 times to remove residual bacteria in the worm pellets. Worm pellets were lyzed by sonication, and the protein concentration of the lysate was determined using the Pierce BCA Protein Assay Kit (Thermo-Scientific). The fatty acid C13:0 (NuChek Prep, dissolved in methanol) was added to each sample to serve as the internal reference control for variations introduced during derivatisation and extraction steps. Fatty acids were derivatised into their respective fatty acid methyl ester (FAME) by incubation in 2% H_2_SO_4_ (Sigma Aldrich) in methanol (Fisher) at 55°C overnight. The reaction was stopped by the addition of 1.5 ml water (Fisher, MS grade). FAMEs were extracted in a 300 μl hexane (Sigma Aldrich) by vigorous vortexing and centrifuging at 1000 rpm for 1 minute. The hexane layer containing the FAMEs was transferred into an amber GC vial (Agilent). FAME analysis was performed using an Agilent 7890A gas chromatograph equipped with an HP-5MS column. Each FAME peak was identified based on its retention time and unique fragmentation ions and quantified using a serial dilution standard curve. FAME abundance measured by GC-MS was normalised to the internal C13:0 reference control of each sample. For each sample, FAMEs concentration (μg/ml) was normalised to protein concentration (mg/ml) as microgram of fatty acid detected per milligram of protein (μg/mg). The fatty acid concentration (μg/mg) for each mutant was normalised to the fatty acid concentration (μg/mg) of N2 control. The final ratio is expressed as relative fatty acid levels in the graph. Three experiments were carried out with three replicates each. Relative fatty acid abundances were plotted in Prism 8. Statistically, significant differences between samples were assessed using the unpaired, non-parametric Mann Whitney test with Benjamini and Hochberg test for multiple hypothesis correction.

### Western blots

More than 2000 day-one adults were used for each sample. Worms were collected in M9 buffer and washed three times in M9. Liquids were removed after centrifugation, and samples were frozen in liquid nitrogen. For worm lysis, RIPA buffer containing protease inhibitors (Sigma-Aldrich, 8340) and phosphatase inhibitors (Roche, 04906845001) was added to each sample at the same volume as the worm pellet. Worms were lysed via sonication (Qsonica, Q700). Protein concentration was measured using Pierce BCA protein assay kit (Thermo Fisher Scientific, PI23227) following manufacturer’s instructions. 4x Leammli sample buffer (Bio-Rad, 1610747) was added to denature the proteins, and samples were heated to 95 °C for 5 min. Samples containing 30-40 mg protein were loaded to 10-20% Tris-Glycine gels (Thermo Fisher Scientific XP10205BOX) for SDS-PAGE. Proteins were transferred to PVDF membranes (Thermo Fisher Scientific, LC2005) and blocked with 5% non-fat milk powder in TBST. The following primary antibodies were used at the indicated dilutions: rabbit α-H3K4me3 (1:1000, Abcam ab8580), rabbit α-H3K9ac (1:1000, Abcam ab10812), rabbit α-H3K18ac (1:1000, Sigma-Aldrich 07-354), rabbit α-H3K27ac (1: 1000, Abcam ab4729), and rabbit α-H3 (1:1000, Cell Signaling 9715). The secondary antibody was α-rabbit IgG HRP-linked (1:1000, Cell Signaling 7074). Antibody signals were developed using SuperSignal West Pico PLUS Chemiluminescent Substrate (Thermo Fisher Scientific, 34577), and bands were quantified with Fiji ImageJ.

### Single site analyses

Transcription factor binding sites (TFBS) analysis was done using oPOSSUM program, version 3.0 ^29^. The parameters used in the oPOSSUM were as follows: conservation cut-off = 0.4, matrix score threshold = 85%, analyzed sequences = 1 kb upstream of transcription start sites, Z-score cut-off = 10, Fisher score cut-off = 7.

### In silico analysis for protein-protein interaction networks

Immunoprecipitation data was used to identify protein-protein interactions for CRTC-1. The analysis was done using STRING, version 11.0 (http://www.string-db.org). The parameters used in the STRING were as follows: network type = full network, meaning of network edges = evidence, active interaction sources = all options (textmining, experiments, databases, co-expression, neighbourhood, gene fusion, and co-occurrence), minimum required interaction score = 0.400, maximum number of interactions to show = no more than 5 interactions (1^st^ shell).

### Tissue-enrichment analysis

Tissue-enrichment analysis was performed using the Enrichment Analysis tool ^66^ from WormBase version WS280. q value threshold = 0.1 was used as parameter.

### Quantification and statistical analysis

Fiji ImageJ was used for image quantifications. GraphPad Prism 9 was used for statistical analyses. Statistical details of experiments can be found in the Figures’ legends. The Log-rank (Mantel-Cox) method was used to compare survival curves. The Mann-Whitney test was used to analyse GFP::SBP-1 and CREp::GFP expressions. Two sample t-test was used to compare IP samples in volcano plot using Perseus software 1.5.1.6. Mann Whitney test with Benjamini and Hochberg test for multiple hypothesis correction was used to compare fatty acid levels. The unpaired t-test was used to compare the expression of antibody signals in immunofluorescence samples and Western blots. For RNA-Seq analysis, differentially expressed genes were identified using the Wald test. For all experiments, P values < 0.05 were considered significant.

## EXTENDED DATA AND SUPPLEMENTARY TABLES TITLES

**Extended Data Fig. 1.**
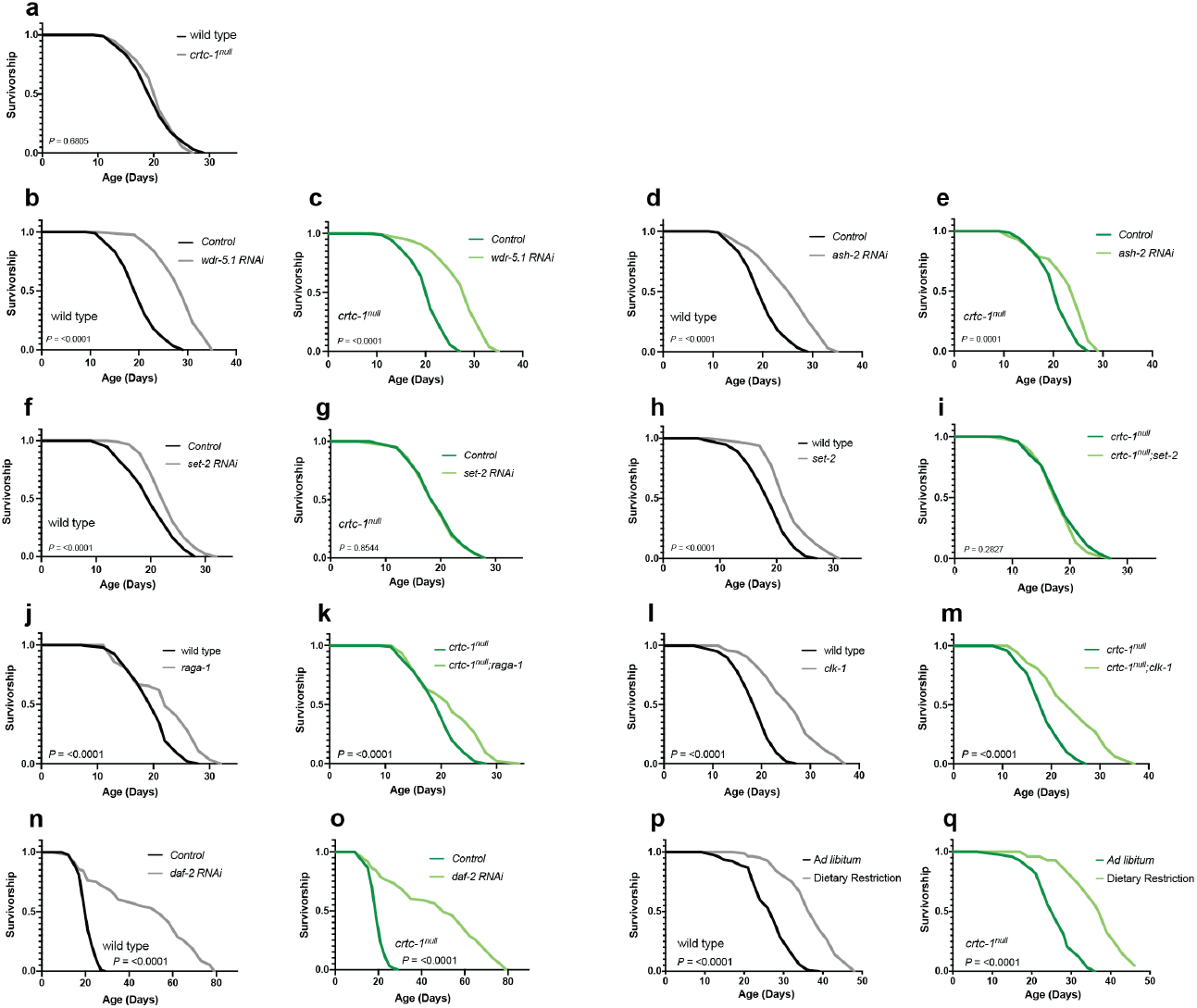
CRTC-1 specifically regulates H3K4me3-mediated longevity. **a**, Lifespan curve showing that *crtc-1^null^* mutant has a similar lifespan to wild-type animals. **b-i**, *crtc-1^null^* mutant has no effect on lifespan extension by *wdr-5.1 RNAi* (**b-c**), reduces *ash-2 RNAi*-mediated longevity (**d-e**), and fully suppresses lifespan extension in *set-2 RNAi* (**f-g**) animals and *set-2(ok952)*mutants (**h-i**). **j-o**, Lifespan curves showing that CRTC-1 is not required to extend lifespan in *raga-1(ok386)* (**j-k**), *clk-1(qm30)* (**l-m**), *daf-2 RNAi* (**n-o**) worms, and dietary restriction (**p-q**). Survival curves compared by the Log-rank (Mantel-Cox) method (see Supplementary Table 2 for lifespan statistics).

**Extended Data Fig. 2.**
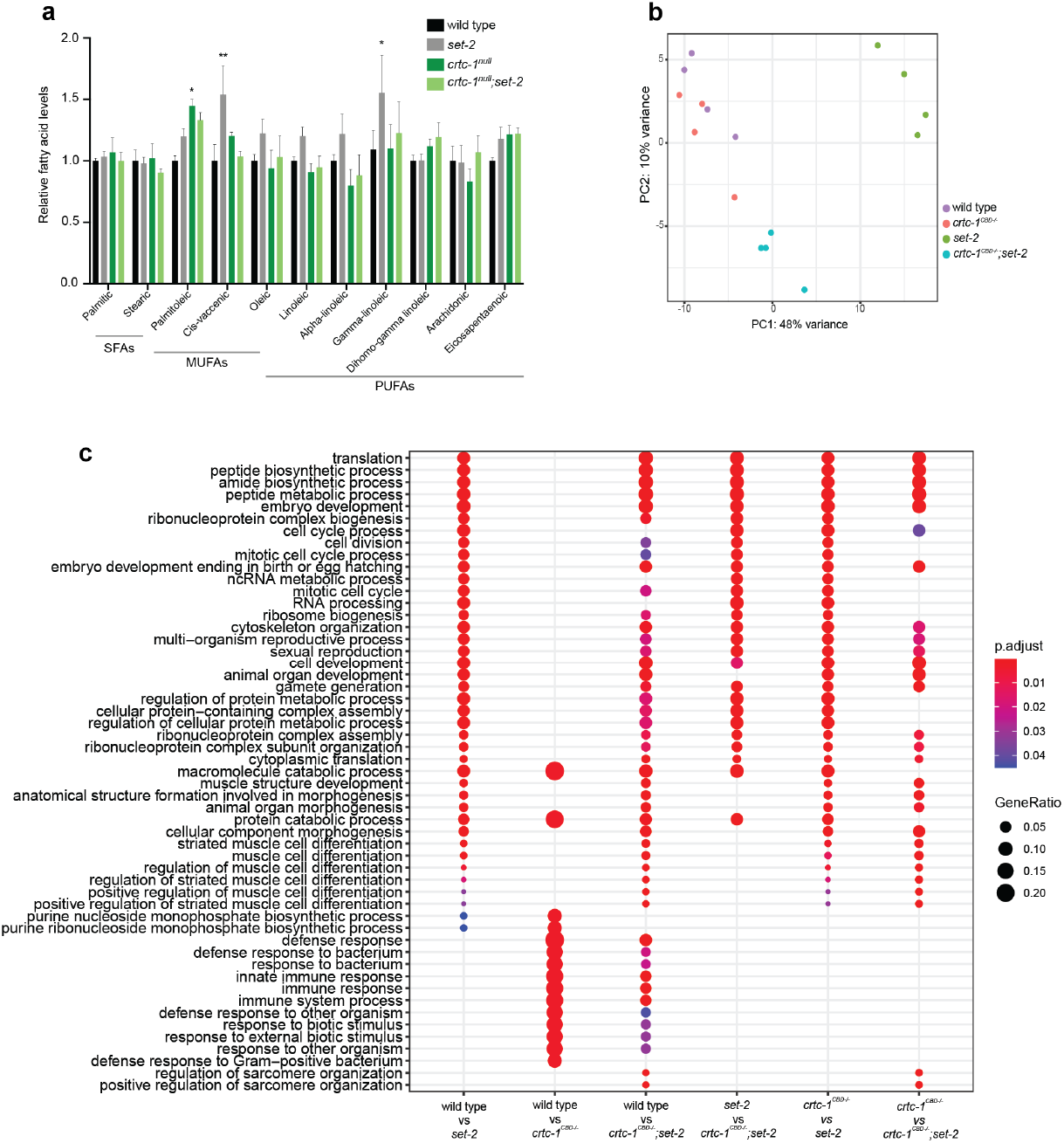
CRTC-1^CBD^ drives a transcriptional shift in H3K4me3-deficient animals. **a**, GC-MS quantification of fatty acids. Mean ± SEM, \**P*<0.05 and \*\**P*<0.01 by unpaired two-way ANOVA. **b**, Principal component analysis of the RNA-Seq data showing similarity between samples. **c**, Over-representation analysis of the Gene Ontology biological process terms for genes comprising all comparisons from the RNA-Seq samples: wild-type, *set-2, crtc-1^CBD-/-^*, and *crtc-1^CBD-/-^;set-2* strains.

**Extended Data Fig. 3.**
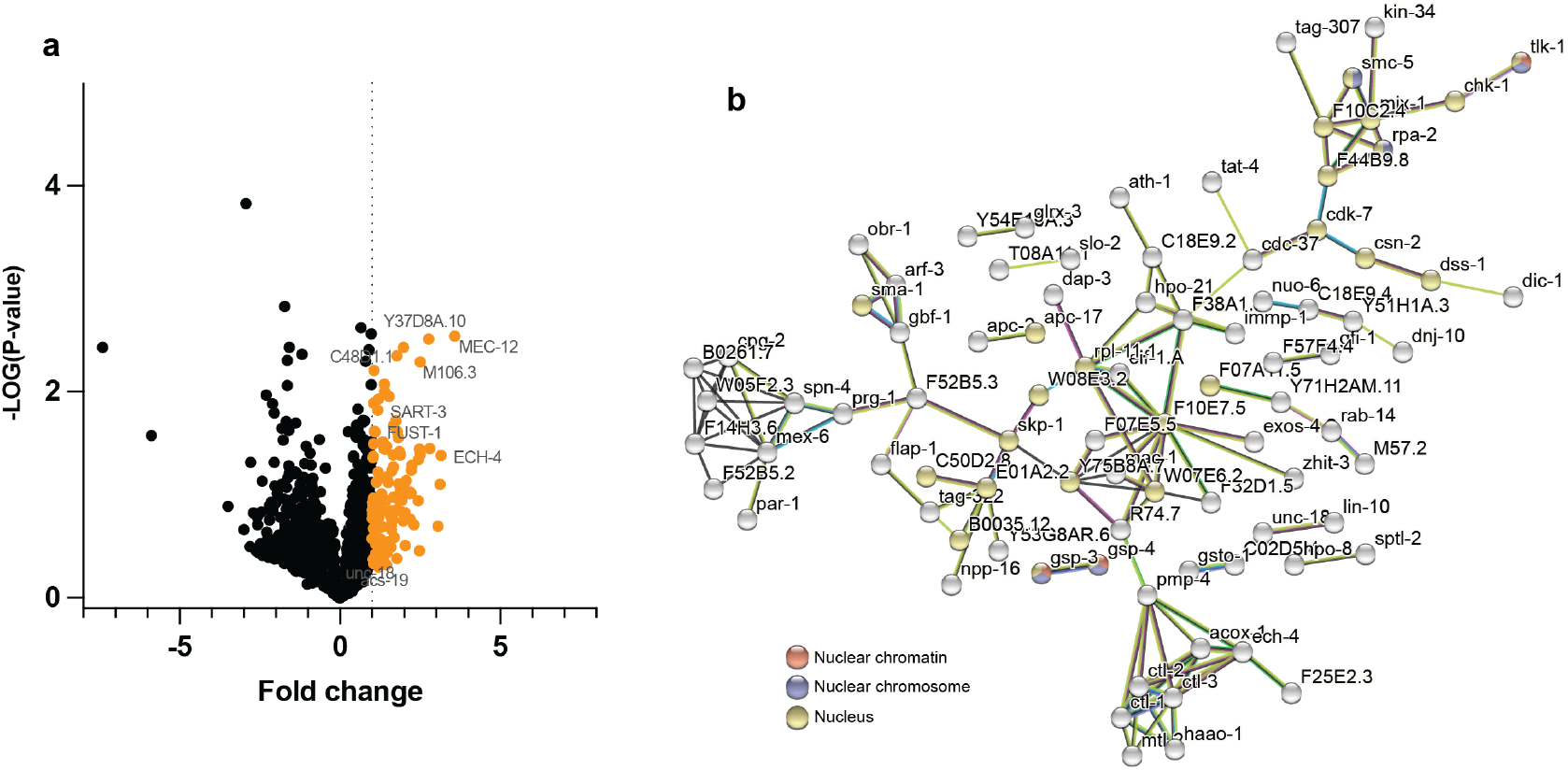
CBD-dependent binding proteins in CRTC-1. **a**, Volcano plot showing proteins identified during immunoprecipitation analysis of endogenously tagged strains CRTC-1::3xFLAG relative to CRTC-1 ^CBD-/-^::3xFLAG strain in the *set-2(ok952)* background. The dotted line indicates a one-fold change. Four biological replicates. *P* values by two-sample t-test. See Supplementary Table 5 for IP statistics. **b**, STRING protein-protein interaction network of CRTC-1 ^CBD^-dependent binding proteins identified during immunoprecipitation analysis. The network nodes are proteins. The edges represent the predicted functional associations. Redline (presence of fusion evidence), green line (neighbourhood evidence), blue line (co-occurrence evidence), purple line (experimental evidence), yellow line (text mining evidence), light blue line (database evidence), and blackline (co-expression evidence).

**Extended Data Fig. 4.**
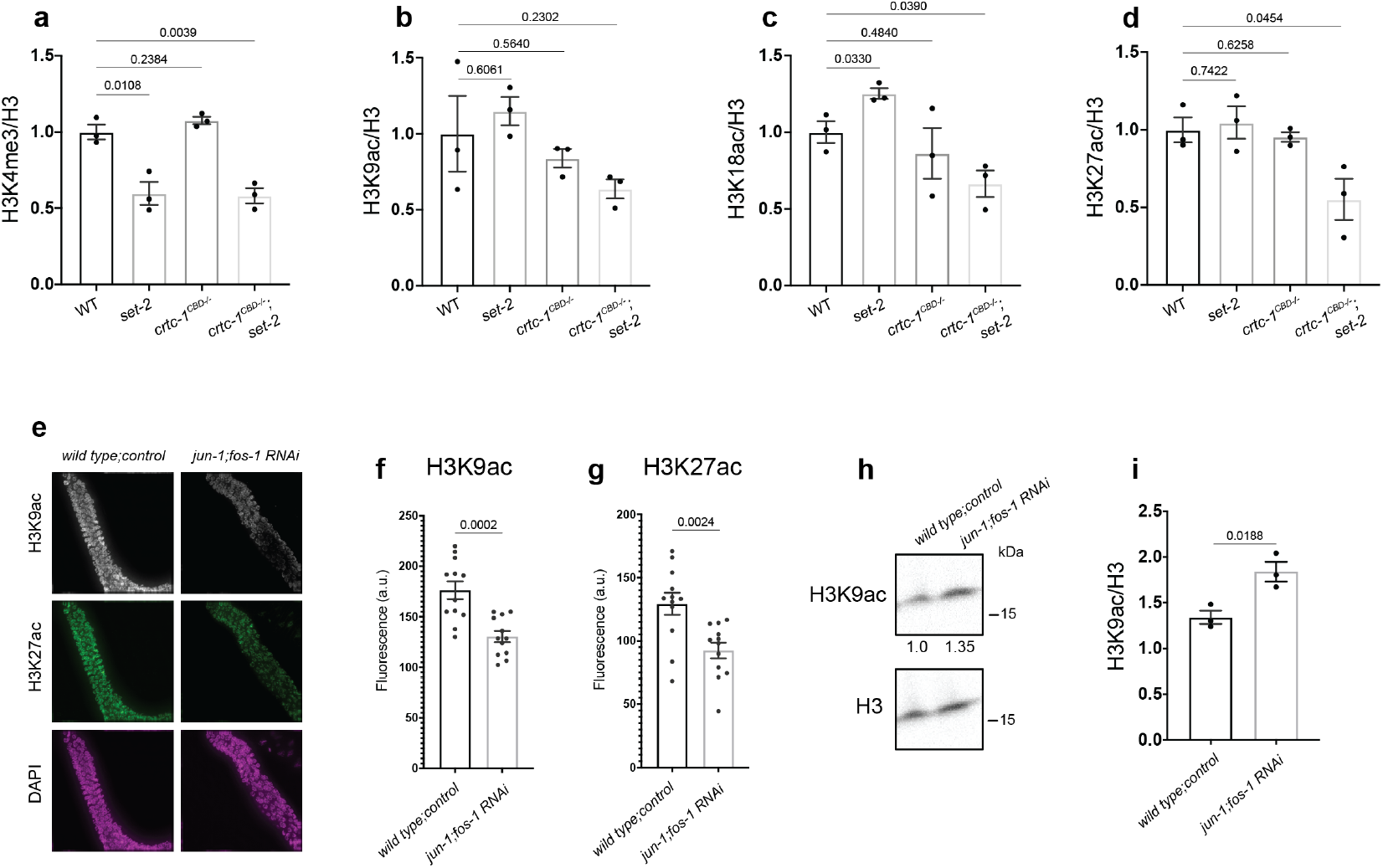
The AP-1 complex regulates levels of histone acetylation. **a-d**, Quantification of Western blots of wild-type, *set-2, crtc-1^CBD-/-^,* and *crtc-1^CBD-/-^;set-2* worms for H3K4me3 (**a**), H3K9ac (**b**), H3K18ac (**c**), and H3K27ac (**d**) marks. H3 used as loading control, and values represent the mean of three independent blots. *P* values by unpaired t-test. **e**, Fluorescence images of the distal region of the gonad from wild-type and *jun-1(gk557);fos-1RNAi* animals co-immunostained for H3K9ac (grey), H3K27ac (green), and DAPI (magenta). **f-g**, Quantification of fluorescence intensity detected for H3K9ac (**f**) and H3K27ac (**g**), indicating a significant reduction in both marks in *jun-1(gk557);fos-1RNAi* animals. Mean ± SEM of n = 10-14 gonads, pooled from three independent experiments. *P* values by unpaired t-test. **h**, Western blot of wild-type and *jun-1(gk557);fos-1RNAi* animals for H3K9ac. **i**, Quantification of Western blots showing an increase of H3K9ac in *jun-1(gk557);fos-1RNAi* animals. H3 used as loading control, and values represent the mean of three independent blots. *P* values by unpaired t-test.

**Extended Data Fig. 5.**
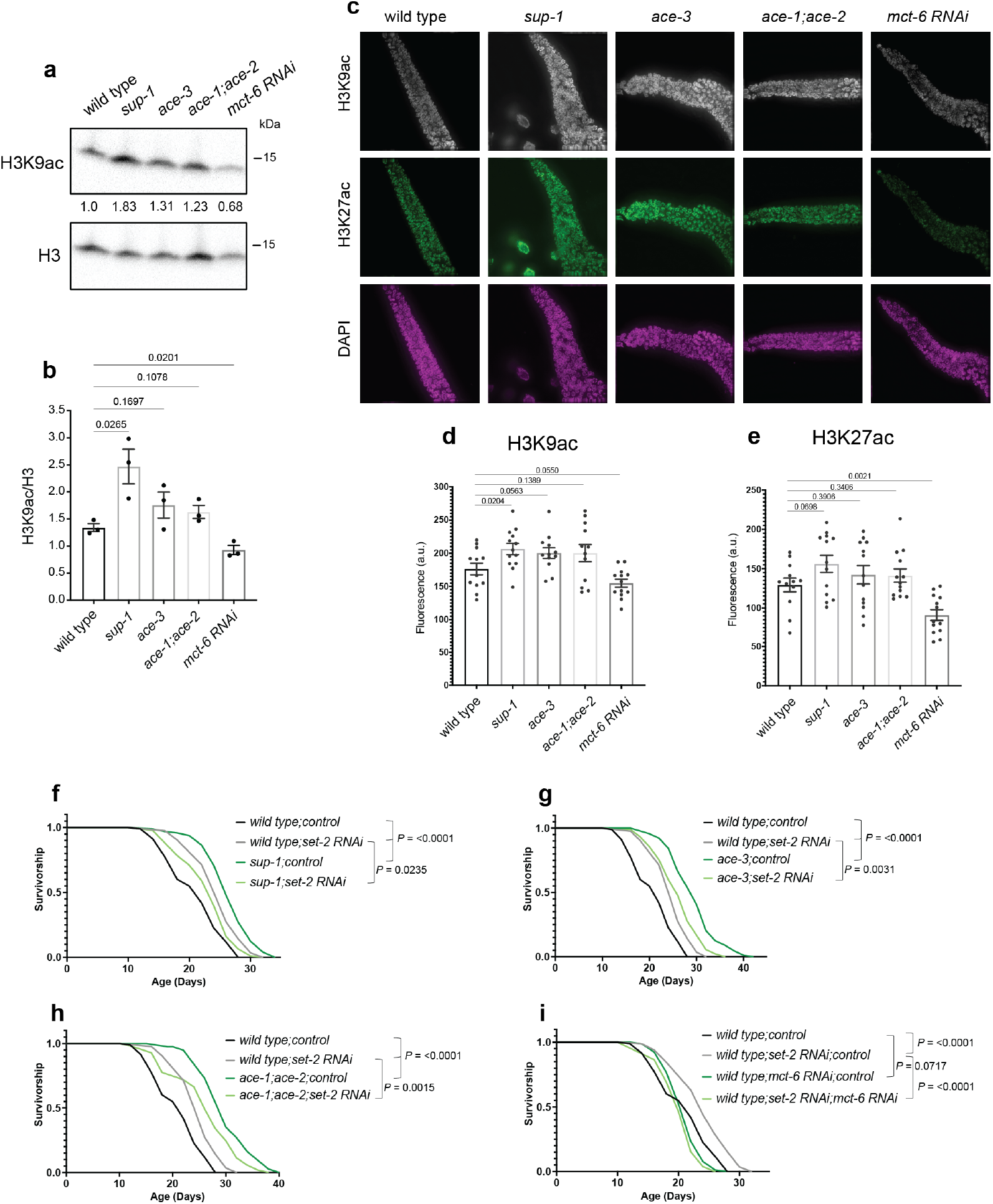
The acetylcholine pathway modulates histone acetylation levels and lifespan extension. **a**, Western blot of wild-type, *sup-1(e995 e2636), ace-3(dc2), ace-1(p1000);ace-2(g72)*, and *mct-6RNAi* animals and the quantification (**b**). H3 levels used as loading control, and values represent the mean of three independent blots. *P* values obtained by unpaired t-test. **c**, Fluorescence images of the distal region of the gonad from wild-type, *sup-1(e995 e2636), ace-3(dc2), ace-1(p1000);ace-2(g72)*, and *mct-6RNAi* animals co-immunostained for H3K9ac (grey), H3K27ac (green), and DAPI (magenta). **d-e**, Quantification of fluorescence intensity detected for H3K9ac (**d**) and H3K27ac (**e**). Mean ± SEM of n = 10-12 gonads, pooled from three independent experiments. *P* values by unpaired t-test. **f-h**, Survival curves showing a lifespan extension in *sup-1(e995 e2636), ace-3(dc2)*, and *ace-1(p1000);ace-2(g72)* mutants but not further extension in lifespan during the *set-2* knockdown. **i**, Survival curves demonstrating that knockdown of *mct-6* blocks the lifespan extension in *set-2RNAi.***j**, Survival curves showing that overexpression of PCAF-1 do not further extend the lifespan in *set-2(ok952)* mutants. Survival curves compared by the Log-rank (Mantel-Cox) method (see Supplementary Table 2 for lifespan statistics).

Supplementary Table 1. List of proteins that interact with CRTC-1.

Supplementary Table 2. Lifespan statistics.

Supplementary Table 3. List of differential gene expression (DGE), gene ontology (GO), and tissue enrichment analyses from RNA-seq.

Supplementary Table 4. List of AP-1 targets in group 1.

Supplementary Table 5. List of proteins that interact with CRTC-1 in a CBD-dependent manner in *set-2* mutants.

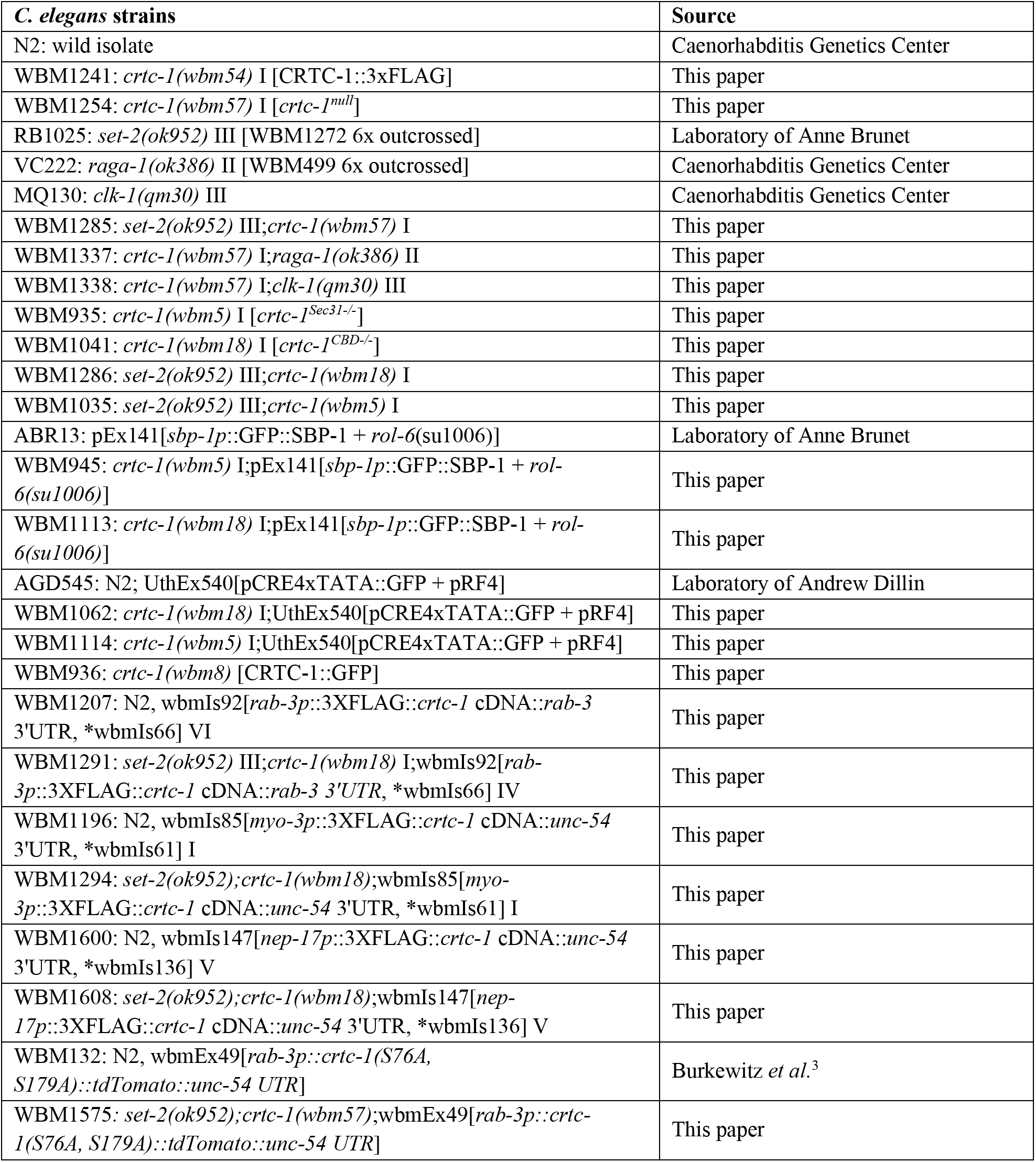
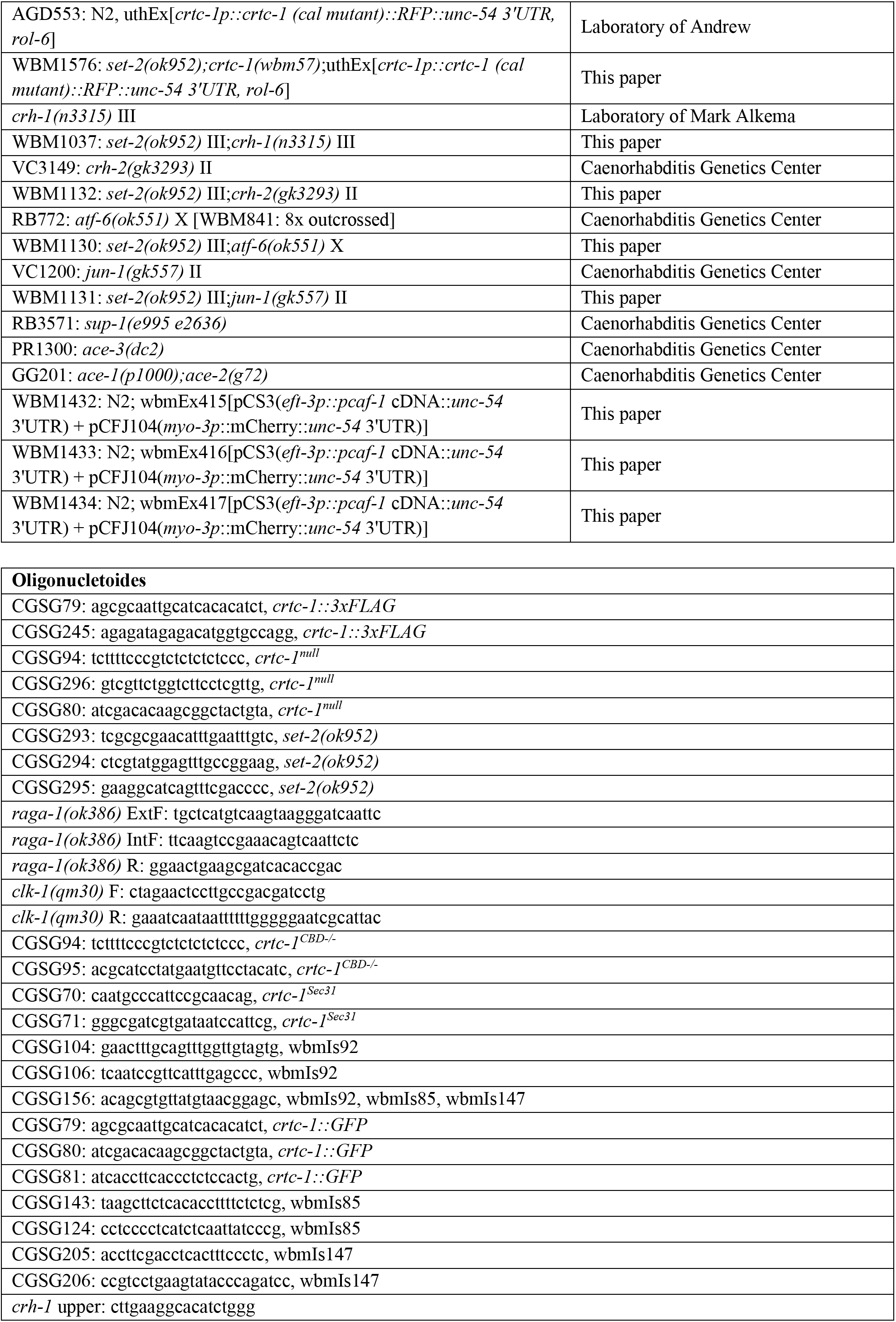
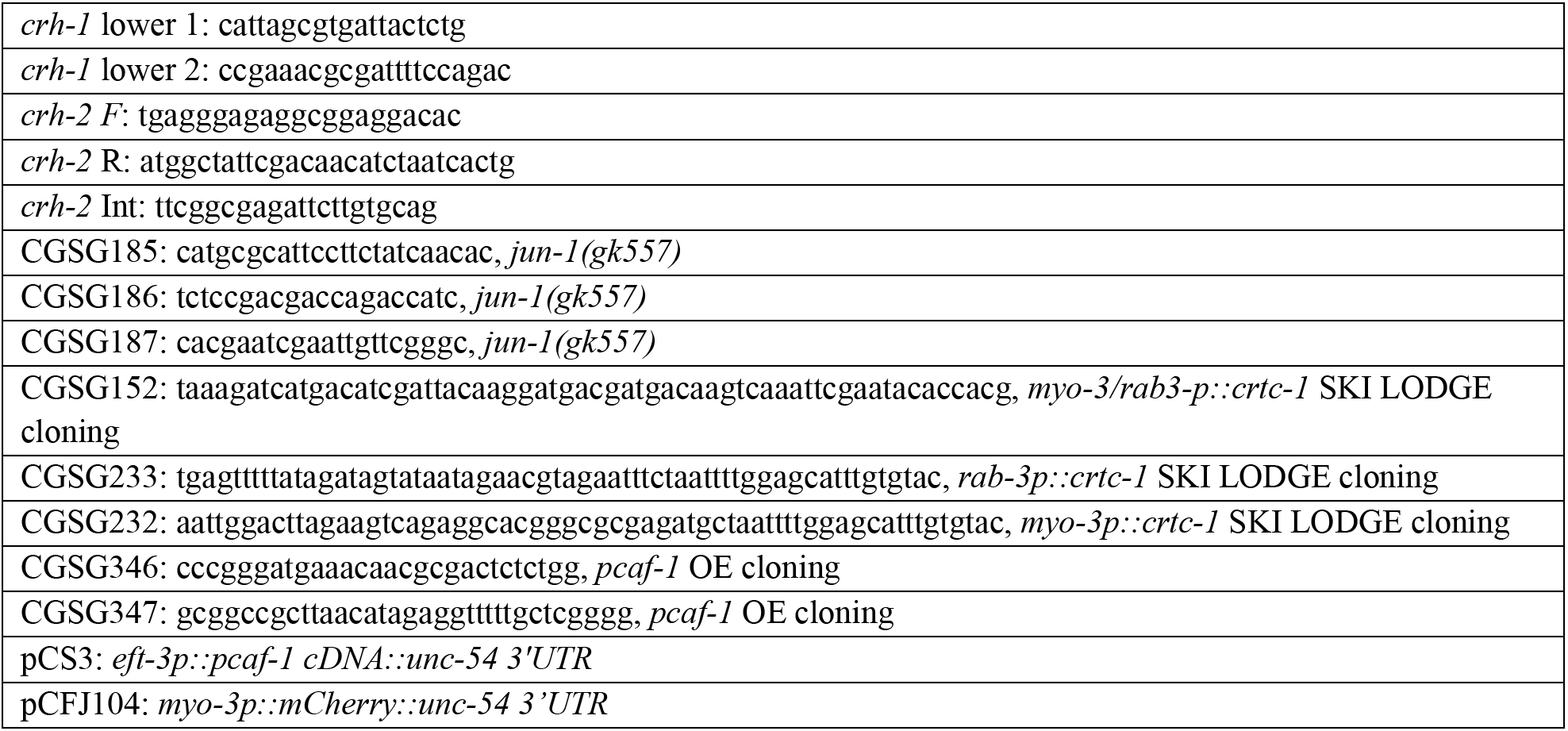

## REFERENCES

1. Corrales, G. M. & Alic, N. Evolutionary Conservation of Transcription Factors Affecting Longevity. Trends Genet 36, 373–382 (2020).

2. Mair, W. et al. Lifespan extension induced by AMPK and calcineurin is mediated by CRTC-1 and CREB. Nature 470, 404–408 (2011).

3. Burkewitz, K. et al. Neuronal CRTC-1 Governs Systemic Mitochondrial Metabolism and Lifespan via a Catecholamine Signal. Cell 160, 842–855 (2015).

4. Ravnskjaer, K. et al. Glucagon regulates gluconeogenesis through KAT2B- and WDR5-mediated epigenetic effects. The Journal of clinical investigation 123, 4318–28 (2013).

5. Liu, Y. et al. A fasting inducible switch modulates gluconeogenesis via activator/coactivator exchange. Nature 456, 269–273 (2008).

6. Han, J. et al. The CREB coactivator CRTC2 controls hepatic lipid metabolism by regulating SREBP1. Nature 524, 243–246 (2015).

7. Amelio, A. L., Caputi, M. & Conkright, M. D. Bipartite functions of the CREB co-activators selectively direct alternative splicing or transcriptional activation. The EMBO Journal 28, 2733–2747 (2009).

8. Escoubas, C. C., Silva-García, C. G. & Mair, W. B. Deregulation of CRTCs in Aging and Age-Related Disease Risk. Trends in Genetics 33, 303–321 (2017).

9. Conkright, M. D. et al. TORCs: transducers of regulated CREB activity. Molecular cell 12, 413–23 (2003).

10. Screaton, R. A. et al. The CREB coactivator TORC2 functions as a calcium- and cAMP-sensitive coincidence detector. Cell 119, 61–74 (2004).

11. Szklarczyk, D. et al. STRING v11: protein–protein association networks with increased coverage, supporting functional discovery in genome-wide experimental datasets. Nucleic Acids Res 47, gky1131 (2018).

12. Greer, E. L. et al. Members of the H3K4 trimethylation complex regulate lifespan in a germline-dependent manner in C. elegans. Nature 466, 383 (2010).

13. Han, S. et al. Mono-unsaturated fatty acids link H3K4me3 modifiers to C. elegans lifespan. Nature 544, 185 (2017).

14. Silva-García, C. G. & Mair, W. B. Confirming the Prolongevity Effects of H3K4me3-deficient set-2 Mutants in Extending Lifespan in C. elegans. Biorxiv 2022.08.02.502497 (2022) doi: 10.1101/2022.08.02.502497.

15. Zhang, Y. et al. Neuronal TORC1 modulates longevity via AMPK and cell nonautonomous regulation of mitochondrial dynamics in C. elegans. Elife 8, e49158 (2019).

16. Kenyon, C., Chang, J., Gensch, E., Rudner, A. & Tabtiang, R. A C. elegans mutant that lives twice as long as wild type. Nature 366, 461–464 (1993).

17. Liu, G. & Sabatini, D. mTOR at the nexus of nutrition, growth, ageing and disease. Nat Rev Mol Cell Bio 21, 183–203 (2020).

18. Dillin, A. et al. Rates of Behavior and Aging Specified by Mitochondrial Function During Development. Science 298, 2398–2401 (2002).

19. Luo, Q. et al. Mechanism of CREB recognition and coactivation by the CREB-regulated transcriptional coactivator CRTC2. Proceedings of the National Academy of Sciences 109, 20865–20870 (2012).

20. Yang, F. et al. An ARC/Mediator subunit required for SREBP control of cholesterol and lipid homeostasis. Nature 442, 700–704 (2006).

21. Hansen, M. et al. Lifespan extension by conditions that inhibit translation in Caenorhabditis elegans. Aging Cell 6, 95–110 (2007).

22. Pan, K. Z. et al. Inhibition of mRNA translation extends lifespan in Caenorhabditis elegans. Aging Cell 6, 111–119 (2007).

23. Troemel, E. R. et al. p38 MAPK Regulates Expression of Immune Response Genes and Contributes to Longevity in C. elegans. Plos Genet 2, e183 (2006).

24. Iourgenko, V. et al. Identification of a family of cAMP response element-binding protein coactivators by genomescale functional analysis in mammalian cells. Proceedings of the National Academy of Sciences of the United States of America 100, 12147–52 (2003).

25. Canettieri, G. et al. The coactivator CRTC1 promotes cell proliferation and transformation via AP-1. Proceedings of the National Academy of Sciences 106, 1445–1450 (2009).

26. Wang, Y., Vera, L., Fischer, W. H. & Montminy, M. The CREB coactivator CRTC2 links hepatic ER stress and fasting gluconeogenesis. Nature 460, 534–537 (2009).

27. Kwon, A. T., Arenillas, D. J., Hunt, R. W. & Wasserman, W. W. oPOSSUM-3: Advanced Analysis of Regulatory Motif Over-Representation Across Genes or ChIP-Seq Datasets. G3 Genes Genomes Genetics 2, 987–1002 (2012).

28. Sui, S. J. H., Fulton, D. L., Arenillas, D. J., Kwon, A. T. & Wasserman, W. W. oPOSSUM: integrated tools for analysis of regulatory motif over-representation. Nucleic Acids Res 35, W245–W252 (2007).

29. Sui, S. J. H. et al. oPOSSUM: identification of over-represented transcription factor binding sites in coexpressed genes. Nucleic Acids Res 33, 3154–3164 (2005).

30. Burkewitz, K. et al. Atf-6 Regulates Lifespan through ER-Mitochondrial Calcium Homeostasis. Cell Reports 32, 108125 (2020).

31. Chinenov, Y. & Kerppola, T. K. Close encounters of many kinds: Fos-Jun interactions that mediate transcription regulatory specificity. Oncogene 20, 2438–2452 (2001).

32. Silva-García, C. G. et al. Single-Copy Knock-In Loci for Defined Gene Expression in Caenorhabditis elegans. G3: Genes, Genomes, Genetics 9, g3.400314.2019 (2019).

33. Li, T. Y. et al. The transcriptional coactivator CBP/p300 is an evolutionarily conserved node that promotes longevity in response to mitochondrial stress. Nat Aging 1, 165–178 (2021).

34. Cohen, E. et al. Caenorhabditis elegans nicotinic acetylcholine receptors are required for nociception. Mol Cell Neurosci 59, 85–96 (2014).

35. Rand, J. Acetylcholine. Wormbook 1–21 (2007) doi:10.1895/wormbook.1.131.1.

36. Mathews, E. A., Mullen, G. P., Hodgkin, J., Duerr, J. S. & Rand, J. B. Genetic Interactions Between UNC-17/VAChT and a Novel Transmembrane Protein in Caenorhabditis elegans. Genetics 192, 1315–1325 (2012).

37. Bradshaw, P. C. Acetyl-CoA Metabolism and Histone Acetylation in the Regulation of Aging and Lifespan. Antioxidants 10, 572 (2021).

38. Halestrap, A. P. The monocarboxylate transporter family— Structure and functional characterization. Iubmb Life 64, 1–9 (2012).

39. Tasoulas, J., Rodon, L., Kaye, F. J., Montminy, M. & Amelio, A. L. Adaptive Transcriptional Responses by CRTC Coactivators in Cancer. Trends Cancer 5, 111–127 (2019).

40. Greer, E. L. et al. Transgenerational epigenetic inheritance of longevity in Caenorhabditis elegans. Nature 479, 365 (2011).

41. Caron, M. et al. Loss of SET1/COMPASS methyltransferase activity reduces lifespan and fertility in Caenorhabditis elegans. (2021) doi:10.1101/2021.06.07.447374.

42. Giblin, W., Skinner, M. E. & Lombard, D. B. Sirtuins: guardians of mammalian healthspan. Trends Genet 30, 271–286 (2014).

43. Kaeberlein, M., McVey, M. & Guarente, L. The SIR2/3/4 complex and SIR2 alone promote longevity in Saccharomyces cerevisiae by two different mechanisms. Gene Dev 13, 2570–2580 (1999).

44. Rogina, B. & Helfand, S. L. Sir2 mediates longevity in the fly through a pathway related to calorie restriction. P Natl Acad Sci Usa 101, 15998–16003 (2004).

45. Tissenbaum, H. A. & Guarente, L. Increased dosage of a sir-2 gene extends lifespan in Caenorhabditis elegans. Nature 410, 227–230 (2001).

46. Kanfi, Y. et al. The sirtuin SIRT6 regulates lifespan in male mice. Nature 483, 218–221 (2012).

47. Ikeda, T., Uno, M., Honjoh, S. & Nishida, E. The MYST family histone acetyltransferase complex regulates stress resistance and longevity through transcriptional control of DAF-16/FOXO transcription factors. Embo Rep 18, 1716–1726 (2017).

48. Yu, R. et al. Inactivating histone deacetylase HDA promotes longevity by mobilizing trehalose metabolism. Nat Commun 12, 1981 (2021).

49. Wang, Y.-L., Faiola, F., Xu, M., Pan, S. & Martinez, E. Human ATAC Is a GCN5/PCAF-containing Acetylase Complex with a Novel NC2-like Histone Fold Module That Interacts with the TATA-binding Protein*. J Biol Chem 283, 33808–33815 (2008).

50. Beurton, F. et al. Physical and functional interaction between SET1/COMPASS complex component CFP-1 and a Sin3S HDAC complex in C. elegans. Nucleic Acids Res 47, 11164–11180 (2019).

51. Cai, Y. et al. Subunit Composition and Substrate Specificity of a MOF-containing Histone Acetyltransferase Distinct from the Male-specific Lethal (MSL) Complex*. J Biol Chem 285, 4268–4272 (2010).

52. Lee, T. W., David, H. S., Engstrom, A. K., Carpenter, B. S. & Katz, D. J. Repressive H3K9me2 protects lifespan against the transgenerational burden of COMPASS activity in C. elegans. Elife 8, e48498 (2019).

53. Boon, R., Silveira, G. G. & Mostoslavsky, R. Nuclear metabolism and the regulation of the epigenome. Nat Metabolism 1–14 (2020) doi:10.1038/s42255-020-00285-4.

54. Paix, A., Folkmann, A., Rasoloson, D. & Seydoux, G. High Efficiency, Homology-Directed Genome Editing in Caenorhabditis elegans Using CRISPR-Cas9 Ribonucleoprotein Complexes. Genetics 201, 47–54 (2015).

55. Evans, T. Transformation and microinjection. WormBook (2006) doi:10.1895/wormbook.1.108.1.

56. Weir, H. J. et al. Dietary Restriction and AMPK Increase Lifespan via Mitochondrial Network and Peroxisome Remodeling. Cell Metab 26, 884–896.e5 (2017).

57. Dobin, A. et al. STAR: ultrafast universal RNA-seq aligner. Bioinformatics 29, 15–21 (2013).

58. Patro, R., Duggal, G., Love, M. I., Irizarry, R. A. & Kingsford, C. Salmon provides fast and bias-aware quantification of transcript expression. Nat Methods 14, 417–419 (2017).

59. Soneson, C., Love, M. I. & Robinson, M. D. Differential analyses for RNA-seq: transcript-level estimates improve gene-level inferences. F1000research 4, 1521 (2015).

60. Love, M. I., Huber, W. & Anders, S. Moderated estimation of fold change and dispersion for RNA-seq data with DESeq2. Genome Biol 15, 550 (2014).

61. Yu, G., Wang, L.-G., Han, Y. & He, Q.-Y. clusterProfiler: an R Package for Comparing Biological Themes Among Gene Clusters. Omics J Integr Biology 16, 284–287 (2012).

62. Lascarez-Lagunas, L. I., Herruzo, E., Grishok, A., San-Segundo, P. A. & Colaiácovo, M. P. DOT-1.1-dependent H3K79 methylation promotes normal meiotic progression and meiotic checkpoint function in C. elegans. Plos Genet 16, e1009171 (2020).

63. Rappsilber, J., Ishihama, Y. & Mann, M. Stop and Go Extraction Tips for Matrix-Assisted Laser Desorption/Ionization, Nanoelectrospray, and LC/MS Sample Pretreatment in Proteomics. Anal Chem 75, 663–70 (2003).

64. Cox, J. & Mann, M. MaxQuant enables high peptide identification rates, individualized p.p.b.-range mass accuracies and proteome-wide protein quantification. Nat Biotechnol 26, 1367–1372 (2008).

65. Cox, J. et al. Andromeda: A Peptide Search Engine Integrated into the MaxQuant Environment. J Proteome Res 10, 1794–1805 (2011).

66. Angeles-Albores, D., Lee, R. Y. N., Chan, J. & Sternberg, P. W. Tissue enrichment analysis for C. elegans genomics. Bmc Bioinformatics 17, 366 (2016).

